# Sex-dependent effects of maternal high-fat diet during lactation in adult THY-Tau22 mice offspring

**DOI:** 10.1101/2025.06.10.658847

**Authors:** Thibaut Gauvrit, Hamza Benderradji, Alexandre Pelletier, Kévin Carvalho, Soulaimane Aboulouard, Emilie Faivre, Estelle Chatelain, Hugo Cannafarina, Léna Labous, Agathe Launay, Marie Fourcot, Dimitri Kwiatkowski, Léna Chesnais, Emmanuelle Vallez, Tristan Cardon, Aude Deleau, Bryan Thiroux, Sabiha Eddarkaoui, Anna Bogdanova, Mélanie Besegher, Fabien Delahaye, Jean-Sébastien Annicotte, Stéphanie Le Gras, Anne Tailleux, Michel Salzet, Guillemette Marot, Luc Buée, David Blum, Didier Vieau

## Abstract

The perinatal environment has been suggested to participate to the development of tauopathies and Alzheimer’s disease but the molecular and cellular mechanisms involved remain contradictory and under-investigated. Here, we evaluated the effects of a maternal high-fat diet (HFD) during lactation on the development of tauopathy in the THY-Tau22 mouse strain, a model of progressive tau pathology associated with cognitive decline.

During lactation, dams were fed either a chow diet (13.6% of fat) or a HFD (58% of fat). At weaning, offspring was fed a chow diet until sacrifice at 4 months of age (the onset of tau pathology) or 7 months of age (the onset of cognitive impairment).

During lactation, maternal HFD increased body weight gain in offspring. At 3 months of age, maternal HFD led to a mild glucose intolerance only in male offspring. Moreover, it impaired spatial memory in both male and female 6-month-old offspring, with males being more impacted. These cognitive deficits were associated with increased phosphorylation of hippocampal tau protein-observed at 4 months in males and at 7 months in females, highlighting a sex-specific temporal shift. Additionally, maternal HFD modified adult hippocampal neurogenesis (AHN), leading to an increase of mature neuronal cells number in females and of dendritic arborization length in males. Synaptic analysis further revealed that maternal HFD led to synaptic loss only in males. Finally, multi-omics approaches showed that maternal HFD has long-term consequences on both transcriptome, proteome and regulome, this effect being also sex-dependent with mitochondrial pathways, ribosomal activity, cilium and the extracellular matrix predominantly impacted in males, while gliogenesis, myelination and synaptic plasticity were primarily affected in females. Regulome analysis suggested that this sex-dependent phenotype was more related to a temporal shift rather than distinct sex-specific alterations. Collectively, our data suggest that maternal malnutrition accelerates the development of tauopathy in THY-Tau22 offspring, with sex-dependent effects, males being impacted earlier than females. These findings highlight the critical role of the perinatal environment as a key window of opportunity for interventions aimed at preventing the development of neurodegenerative diseases.

## Introduction

Alzheimer’s disease (AD) is the most common form of dementia.^1^ It is characterised by the presence of two lesions in the brain leading to the clinical symptoms: amyloid deposits composed of aggregated beta amyloid peptides and neurofibrillary tangles (NFT) which correspond to the accumulation of misfolded hyper- and abnormally phosphorylated tau proteins.^2–4^ Besides AD, such tau pathology, also known as tauopathy, is a common feature of several other neurodegenerative diseases such as progressive supranuclear palsy, Pick’s disease or corticobasal degeneration.^5,6^ In AD patients, the spatiotemporal evolution of misfolded tau from entorhinal cortex to hippocampus and finally neocortex is closely correlated with cognitive deficits, supporting a key role of tau-related mechanisms.^7,8^

AD is mainly multifactorial, combining genetic and environmental risk factors, the main cause being ageing.^9,10^ As women have a longer life expectancy, they are also more at risk of developing the disease, but the biological explanations remain poorly understood.^11^ Epidemiological studies reveal that cardiometabolic disorders,^12,13^ type 2 diabetes^14–16^ or obesity^17,18^ increase the risk to develop AD, while Mediterranean diet,^19,20^ moderate caffeine consumption^21,22^ or physical activity^23^ are beneficial. These population studies were corroborated by experimental studies in models.^24–29^ For instance, long-term high-fat diet (HFD) in adult AD mouse models has been shown to worsen neuropathological lesions and/or memory impairments.^25,26,30,31^ Essentially, all these studies evaluated the impact of these environmental factors in the adulthood or the elderly. In sharp contrast, only few studies focused on earlier stages such as perinatal life (gestation and/or lactation), infancy and adolescence although these periods are crucially sensitive to environmental factors.^32^ Indeed, it is now widely accepted that environmental impairments, such as malnutrition, during the early stages of development (gestation and/or lactation) can have long-lasting detrimental impacts and increase the risk to develop multiple disorders in offspring. This concept of perinatal programming is known as “developmental origin of health and diseases” (DOHaD) or Barker’s hypothesis.^33–35^ Epidemiological data have particularly reported that maternal obesity increases the risk of developing metabolic and cognitive disorders in offspring during adolescence.^36–39^ A growing body of evidence in non-transgenic animals has also revealed that a maternal HFD during gestation and/or lactation induces memory impairments in adult offspring.^37,40–44^ These observations were ascribed to synaptic alterations,^41,42,44–48^ neurogenesis impairments,^44,49–51^ neuroinflammation^52,53^ and the presence of AD-related changes in the hippocampus of the offspring.^54^ Moreover, we recently reported that maternal HFD only during lactation is sufficient to lead to profound modifications of the hippocampal proteome and transcriptome in the offspring in a sex-dependent manner.^44^ Although these works in non-transgenic animals strongly suggest that maternal malnutrition is prone to accelerate the progression of aging-related disorders, very scarce studies are available in mouse models of AD. In the homozygous 3xTgAD mice that develop early amyloid deposits and late tau pathology,^55^ it has been reported that maternal HFD during gestation and lactation programs memory disorders associated with increased AD hallmarks in the adult offspring hippocampus.^56,57^ Puzzlingly, application of a maternal HFD only during gestation was found to reduce cognitive decline as well as amyloid and tau load in adult offspring in the same 3xTgAD strain.^58^ These apparently contradictory results indicate that further work is necessary to determine the underpinning biological mechanisms of perinatal programming of “AD-like pathology” but also suggest that lactation is a very sensitive and suitable period to analyse in order to address the long-term deleterious consequences of a maternal HFD in adult mouse offspring. In rodents, lactation, which corresponds to the third trimester of gestation in human, coincides with the peak of brain development, during which neuronal maturation, dendritic growth, gliogenesis and synaptogenesis take place.^32^

In the present work, we explored the effects of a maternal HFD during lactation on both cognition and hippocampal pathological hallmarks (tau pathology, synapses, neurogenesis, neuroinflammation, metabolism) in offspring of both sexes in a model of AD-like tauopathy (THY-Tau22). THY-Tau22 mice develop an early and progressive tau pathology in the hippocampus associated with memory deficits, underlined by synaptic impairments.^59–62^ To uncover the cellular and molecular hippocampal impacts of maternal HFD in THY-Tau22 offspring, we have carried out unsupervised approaches using a multi-omic strategy (transcriptomic, proteomic and regulomic). Our data demonstrate that maternal HFD during lactation leads to worsening of tau pathology, synaptic loss and neurogenesis impairments in the hippocampus of adult offspring in a sex-dependent manner, converging towards spatial memory worsening. In addition, we show that maternal HFD has sex-dependent long-term consequences on both transcriptome and proteome with mitochondrial pathways, ribosomal activity, cilium and the extracellular matrix predominantly impacted in males, while gliogenesis, myelination and synaptic plasticity related pathways were rather affected in females. These data support the idea that age-related neurodegenerative diseases may have, at least in part, a neurodevelopmental origin.

## Materials & Methods

### Experimental protocol

Heterozygous THY-Tau22 mice of C57Bl6/J genetic background overexpress the human mutated (G472V and P301S) 1N4R tau protein isoform under the control of the Thy1.2 neuronal promotor. THY-Tau22 mice develop a progressive hippocampal tau pathology (from 3 months of age) and spatial memory deficit (from 7-8 months of age). Eight-week-old C57BL6/J female mice were mated with THY-Tau22 male mice (two females per male). After 2 weeks, females were housed individually. During lactation (postnatal day P0 to P21), females were randomly assigned to either a high-fat diet (HFD; 58% of fat, Research Diets D12331) or a control chow diet (13.5% of fat, Safe A03), generating four groups in the offspring: THY-Tau22 Chow (TC) and THY-Tau 22 HFD (TH) mice, and their littermate WT Chow (WC) and WT HFD (WH) mice. Nutrient composition of each diet is given in Supplementary Table 1. At P1, litter size was adjusted to six pups per dam. At weaning (P21), offspring returned to a standard diet (8.4% of fat, Safe A04) and housed according to sex and maternal diet (three animals per cage) until the sacrifice either at 4 or 7 months of age (Supplementary Fig. 1). In each group, body weight, food and water intake were recorded weekly. All animals were housed in a pathogen-free facility and maintained under conditions controlled for temperature (22°C and 12-h light/12-h dark cycle) with *ad libitum* access to food and water. Protocols were approved by an ethics committee (protocol #8677-2017011213553351v3).

### Behavioural studies

Behavioural experiments were conducted on 6-month-old mice, randomly assigned, by experimenter blinded to the genotype and phenotype of mice. To allow for habituation, home cages were placed in the dedicated room 30 minutes prior to testing. Each equipment was cleaned with 70% ethanol between animals. Procedures are given in the Supplementary material.

### Sacrifice and tissue dissection

Mice were sacrificed at 4 or 7 months of age, after 6 h of fasting. 4-month-old animals and one part of 7-month-old animals were sacrificed by cervical dislocation, hippocampi were removed and frozen for molecular biology and biochemical analysis. The other part of 7-month-old animals was anesthetized, perfused with saline solution 0.9% and paraformaldehyde 4%, brains were removed, post-fixed in paraformaldehyde 4% (24 h), incubated in a sucrose saline solution (30%) and finally frozen in 2-methyl-butane for immunostaining analysis. Moreover, among these 7-month-old mice, a subgroup of animals was injected beforehand with 5-ethynyl-2’-deoxyuridine (EDU; Sigma-Aldrich 900584) to study hippocampal adult neurogenesis. The hemispheres of these brains were separated: one was incubated in sucrose, frozen, and then cut using a cryostat, while the other was incubated in saline solution with 0.2% of azide before being cut with a vibratome (three-dimensional analysis).

### Metabolic analysis

Blood glucose measurements, glucose tolerance test and biochemical plasma parameters analysis were performed after 6 h of fasting in 4-month-old animals. Procedures are described in the Supplementary material.

### Hippocampal adult neurogenesis analysis

#### 5-ethynyl-2’-deoxyuridine injection

To study the neurogenesis, 6-month-old mice were injected with 5-ethynyl-2’-deoxyuridine (EDU; Sigma-Aldrich 900584), a modified thymidine analogue that incorporates into DNA during cell division. Fifty mg/kg of body weight of EDU were injected 2 times per day for 3 days, 21 days before the sacrifice.

#### 5-ethynyl-2’-deoxyuridine staining

Frozen brains from 7-month-old mice injected with EDU were cut to obtain 35 µm-thick coronal floating sections using a cryostat. EDU was labelled with a fluorochrome using the Click-iT EDU 647 kit and following the manufacturer’s instructions (Sigma-Aldrich BCK-EDU647). Images were taken using a slide scanner (Zeiss Axioscan Z1) with a 20X objective. The number of EDU positive cells in the subgranular zone of the dentate gyrus was manually counted on serial coronal sections representing the entire hippocampus.

### Molecular biology analysis

#### RNA extraction and quantitative PCR analysis

Total RNA was extracted from hippocampi and purified using RNeasy Lipid Tissue Mini kit (QIAGEN 1023539) as described in Gauvrit *et al*.^44^ Quantitative PCR analysis is described in Supplementary material.

#### RNA sequencing

RNA-seq was performed from 300 ng of total RNA (five mice per group, except for 4-month-old male mice, which has four animals) using the same protocol described in Gauvrit *et al*.^44^ Differential analyses of RNA-seq data were performed using Wald tests from the R Bioconductor package DESeq2 (v 1.44.0). Log2 fold changes were shrinked using the lfcShrink function of DESeq2 package. Differences were considered statistically significant when *P* values adjusted for multiple testing using Benjamini-Hochberg correction were less than 0.05 and the absolute value of shrinked log2 fold change was greater than 0.26. Over representation analyses (ORA) were performed using enricher function from the R Bioconductor package clusterProfiler (v 4.12.6), based on Kyoto encyclopedia of genes and genomes (KEGG) and MSigDB gene ontology (GO) pathways, using as background genes present in the considered database. The *P* values of enriched pathways were adjusted using Benjamini-Hochberg procedure, and an adjusted *P* value < 0.05 was considered significant. It is important to note that, while all data were analyzed together, RNA-seq experiments for 4- and 7-month-old male animals were not performed simultaneously and that our experimental design does not allow us to definitely distinguish the effect of age from the effect of sequencing date in male offspring. However, many genes known to be modulated during the progression of tau pathology were found to be deregulated, suggesting that impact of this time shift of sequencing is minimal.

### Biochemical analysis

Biochemical procedures are provided in the Supplementary material.

### Immunostaining analysis

Immunostaining procedures are provided in the Supplementary material.

### Synaptic proteins

Images were taken using a LSM980 confocal microscope (Zeiss) with a 63X objective and the Airyscan option. For each section (two per mouse), two images were taken at the level of the molecular layer of the dentate gyrus and two at the level of the stratum radiatum of cornu ammonis (CA1). Labelling was quantified using Imaris software (v 10.1) and the “Spot” function (estimated diameter = 0.300 μm). This function quantifies the number of dots, each corresponding to a pre-or post-synaptic compartment.

### EDU/DCX and EDU/NeuN co-labelling

Images were taken using a LSM980 confocal microscope (Zeiss) with a 40X objective. Quantification of the number of EDU^+^, EDU^+^/NeuN^+^ and EDU^+^/DCX^+^ labelled cells was performed manually with the oculars and limited to the subgranular zone of the dentate gyrus.

### Tri-dimensional analysis of EDU/DCX cells

Images of each EDU^+^/DCX^+^ cell in the subgranular zone of the dentate gyrus were taken using a spinning disk confocal microscope (Zeiss) with a 40X objective. To obtain the entire EDU^+^/DCX^+^ cell (soma and arborization), the z-stack and mosaic options were used. Then, cell reconstruction was performed using the “surface” and “filament” options of the software Imaris (v 10.1) and the distance between the soma and the furthest point reached by the dendritic arborization was calculated.

### Mass spectrometry

A shotgun bottom-up proteomic approach was performed (three mice per group, already used in RNA-seq analysis) as previously described in Gauvrit *et al*.^44^ Differential analyses of proteomics were performed using moderated t-tests from the R Bioconductor package limma (v 3.60.4). Differences were considered statistically significant when *P* values adjusted for multiple testing using Benjamini-Hochberg correction were less than 0.05 and the absolute value of log2 fold change was greater than 0.26. ORA was performed using enricher function from the R Bioconductor package clusterProfiler (v 4.12.6), based on KEGG and MSigDB GO pathways, using as background proteins present in the considered database. The *P* values of enriched pathways were adjusted using Benjamini-Hochberg procedure, and an adjusted *P* value < 0.05 was considered significant.

### Regulome analysis

Regulome analysis correlation of transcription factor (TF) protein abundance and downstream target have been conducted as described in Gauvrit *et al*.^44^ with minor changes. Briefly, this approach uses random forest method based on GENIE3 R package to infer regulatory links between TF and genes^63^ and the CisTarget database (mm10 version 10) to filter for direct links based on TF motif conservation enrichment in transcription start site of the genes.^64^ The minor change is the mice TFs are now annotated thanks to CisTarget database (mm10 v 10), and the TF-gene link score of more than 0.01 was chosen based on knee plot was considered for the subsequent construction of gene modules as described^44^. The following steps of the regulome identification are the same. ORA of each regulon for MSigDB Gene Ontology and Canonical Pathways (c2.cp.v2023 and C5.go.v2023) was conducted using hypergeometric test phyper R function, using as background genes all transcriptionally detected passing quality control (i.e. genes with more than 1 count per million reads in at least 10% of the samples). To assess influence of maternal HFD on regulons, regulons enrichment for genes differentially expressed in maternal HFD vs. Chow was performed using DESeq2 statistics and fgsea package.^65^

### Statistics

Image acquisition and quantification, western blot quantification, as well as metabolic and behavioural evaluations, were performed by investigators blind to the experimental conditions. Values are represented as mean ± standard error of the mean (SEM). Normality of the distribution and homoscedasticity were checked using Shapiro-Wilk test and F or Bartlett tests, respectively. Differences between mean values were determined using two-tailed unpaired Student’s t-test, one sample *t*-test, one-way ANOVA followed by a *post hoc* Tukey’s multiple comparisons test or two-way ANOVA followed by a post hoc Sidak’s multiple comparisons test. To compare proportions, Fisher’s exact test was used. All tests were performed using GraphPad Prism software (v 9.0.0). A *P* value < 0.05 was considered significant.

## Results

### Sex-dependent effects of maternal HFD on physiological and metabolic factors in THY-Tau22 offspring

Maternal HFD decreased the body weight of dams (*P* < 0.001 at P15 and P21, Supplementary Fig. 2A) in spite of increasing food intake (*P* < 0.001 at P21, Supplementary Fig. 2B). It also increased the body weight of male (*P* < 0.001, Supplementary Fig. 2C) and female (*P* < 0.001, Supplementary Fig. 2D) THY-Tau22 offspring at weaning (P21). This effect was transient and was no longer observed in adult animals (Supplementary Table 4). Similarly, there was no change in glycemia and insulinemia after 6 hours of fasting, nor in the free fatty acid plasma level and rectal temperature in 4-month-old offspring. However, the triglyceride (*P* < 0.05) and cholesterol (*P* < 0.01) plasma levels were decreased, respectively in male and female offspring (Supplementary Table 4). Finally, IPGTT analysis showed that male offspring exhibited slight glucose intolerance (*P* < 0.01, Supplementary Fig. 2E) that was not observed in females (Supplementary Fig. 2F), revealing a sex-dependent effect at 3-4 months of age, an early pathological stage in the THY-Tau22 model.

### Sex-dependent effects of maternal HFD on spatial memory in adult THY-Tau22 offspring

To assess the impact of maternal HFD on learning and memory, we tested animals at 6-7 months, when, in THY-Tau22 model, tau pathology is ongoing but memory deficits remain limited.^59^ Using actimetry, no difference in spontaneous locomotor activity was observed with comparable distance moved and velocity (Supplementary Fig. 3A-B and 3D-E) over all groups. Anxiety-like behaviour assessed using elevated plus maze revealed, consistently with previous works,^60^ that both male (*P* < 0.001, Supplementary Fig. 3C) and female (*P* < 0.001, Supplementary Fig. 3F) THY-Tau22 offspring spent more time in the open arms than littermate controls. No effect of the maternal diet was observed. We then evaluated short-term spatial memory using the Y-maze task. During the learning phase, all groups spent similar percentage of time in the familiar vs. the start arms (Fig. 1A and 1B). During the test phase, WC, WH and TC male and female explored significantly more than 50% in the new arm (i.e. above the chance level), indicating proper memory retention (Fig. 1C and 1D). In contrast, TH male and female offspring did not (Fig. 1C and 1D), suggesting that maternal HFD during lactation impaired short-term spatial memory at later stage in THY-Tau22 mice. Finally, long-term spatial memory was tested using the Barnes maze.^66,67^ All groups demonstrated a reduced distance to find the escape box over the four days of trials, highlighting their learning abilities (*P* < 0.001, Fig. 1E and 1F). In the probe test performed 24 hours later, results indicate that WC and WH males (*P* < 0.05 for WC and WH, Fig. 1G) as well as WC and WH females (*P* < 0.01 for WC and *P* < 0.001 for WH, Fig. 1H) spent significantly more than 25% of time in the target quadrant (i.e. above the chance level), indicating proper spatial memory. In line, all mice from these groups explored the cumulative zone (i.e. the area comprising the target hole and the two neighbouring one, Fig. 1I and 1J), which can be considered as a precision parameter. THY-Tau22 male mice did not spend more time in the target quadrant than expected by chance (i.e. > 25%, Fig. 1G and 1H), although TC mice spent a greater of time in this quadrant as compared to TH mice (not significant). Interestingly, in agreement, 78% of TC male mice vs. none of the TH male mice explored the cumulative zone (Fig. 1I), suggesting that maternal HFD impaired long-term spatial memory in THY-Tau22 male offspring. In females, TC and TH mice had no preference for the target quadrant (Fig. 1H) and mice similarly explored the cumulative zone (57% of TC vs. 58% of TH mice, Fig. 1J), suggesting that maternal HFD did not affect long-term memory in THY-Tau22 female offspring. Overall, maternal HFD induced early impairments of short-term spatial memory in both male and female, and only affects long-term memory in male offspring, highlighting sex-dependent consequences with males being more affected at this stage than females.

**Figure 1.**
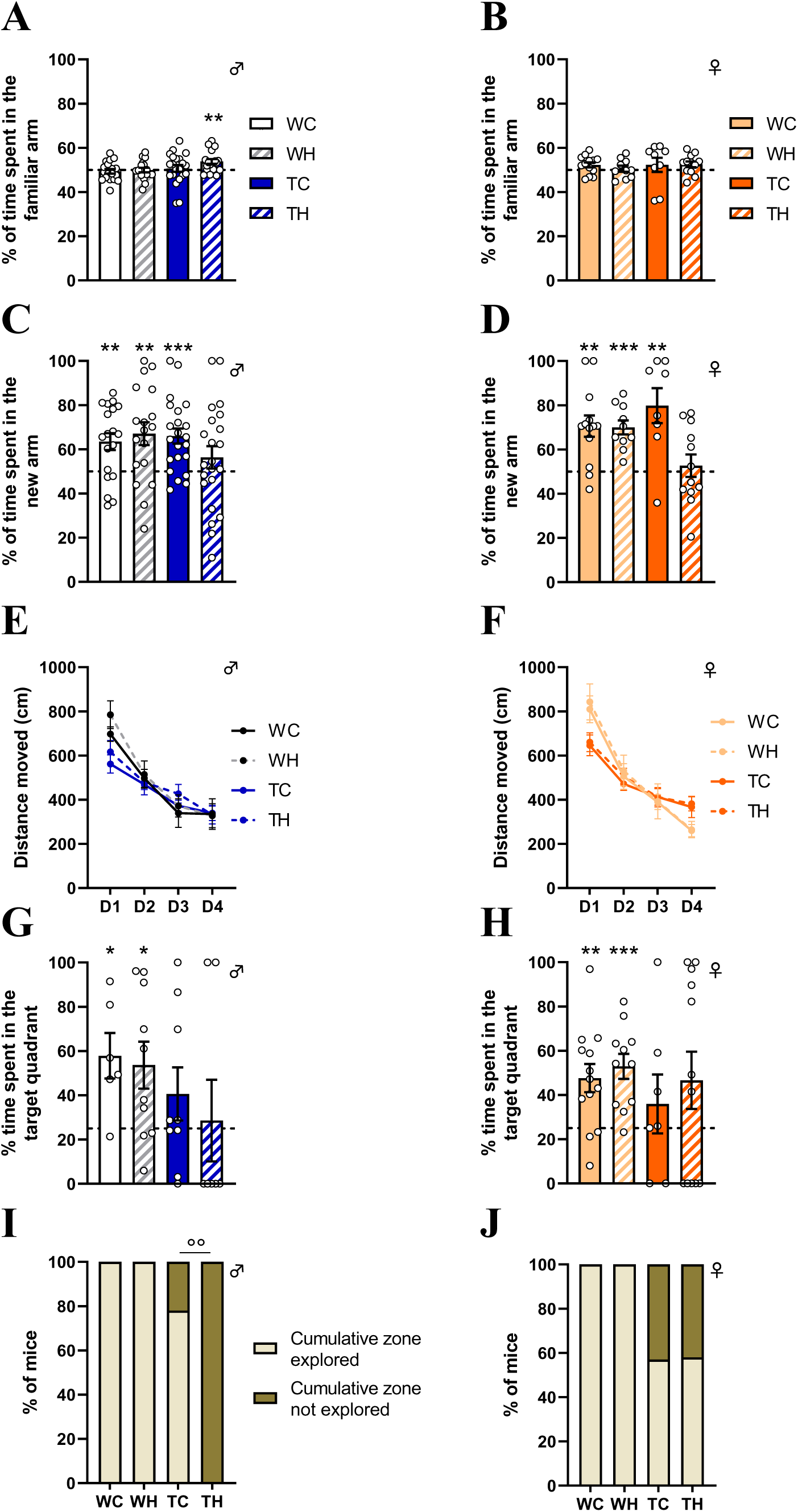
Effect of maternal HFD on spatial memory in adult offspring. Short-term and long-term spatial memory was evaluated in 7-month-old animals using, respectively the Y-maze (**A**-**D**) and the Barnes maze tasks (**E**-**H**). (**A** and **B**) Percentage of time spent in the familiar arm during the learning phase. (**C** and **D**) Percentage of time spent in the new arm during the test phase. (**E** and **F**) Distance moved during the learning phase, representing the animal’s ability to find the escape box during the four days of training. (**G** and **H**) Percentage of time spent in the target quadrant (containing the target hole) during the test phase. (**I** and **J**) Percentage of mice that explored or not the cumulative zone (target hole and the two holes adjacent ones). \*\**P* < 0.01, \*\*\**P* < 0.001 using one sample t-test (theoretical vs. value: 50% for A-D and 25% for G-H). °°*P <* 0.01 using a Fisher’s exact test. *n* = 6-22 per group. Values are expressed as mean ± SEM.

### Sex-dependent effects of maternal HFD on tau pathology in adult THY-Tau22 offspring

At 4 months, maternal HFD significantly increased tau phosphorylation at pY18, pS212/T214 (AT100), pS396 and pS396/S404 in male THY-Tau22 offspring. However, there was a reduction at 7 months at pY18, pT181, pT214, pS396, pS396/S404 and pS422, probably resulting from the increase of total tau protein level (Table 1). Conversely, in female offspring, although the diet had no effect at 4 months, it led to an increase in tau phosphorylation at specific phospho-epitopes (pY18, pT181, pT214, pS212/T214, pS396/S404 and pS422, Table 1) at 7 months. Notably, although a tight link is known between neuroinflammation and tau phosphorylation,^68^ these effects were associated neither with alterations in microglia and astrocytes reactivity (Supplementary Fig. 4), nor major changes of neuroinflammatory markers (Supplementary Fig. 5 and 6). However, in males, *Trem2* (a microglial marker) was slightly and significantly increased by maternal HFD at 4 months (*P* < 0.05), while *Itgax* (another microglial marker) showed a trend to increase at 7 months (*P* = 0.100, Supplementary Fig. 4). Altogether, these data indicate that maternal HFD impacts hippocampal tau pathology, without altering neuroinflammation, in a sex-specific manner, changes being seen earlier in male than in female offspring.

**Table 1.**
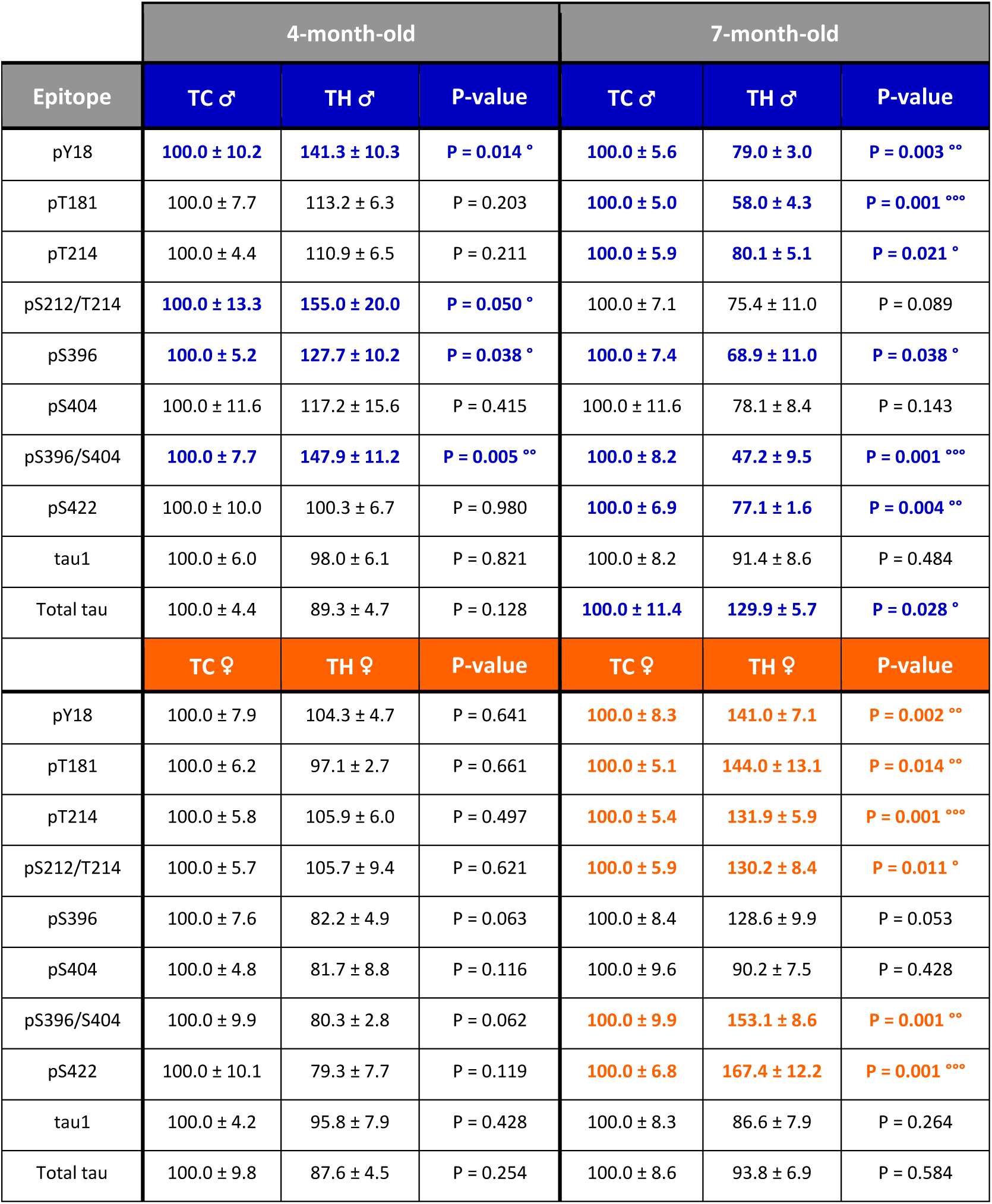
Effect of maternal HFD on hippocampal tau phosphorylation in adult offspring. Western blot quantification of tau phosphorylation at Y18, T181, T214, S212/T214 (AT100), S396, S404, S396/S404 and S422 epitopes, as well as dephosphorylated tau (tau1) in TC and TH mice at 4 or 7 months of age (males and females). Total tau corresponds to the mean of N-terminal and C-terminal tau western blot quantifications. All epitopes were normalized to total tau proteins. °*P* < 0.05, °°*P* < 0.01, °°°*P* < 0.001 versus respective TC mice using Student’s *t*-test. *n* = 8-9 per group. Values are expressed as mean ± SEM.

|  | 4-month-old |  |  | 7-month-old |  |  |
| --- | --- | --- | --- | --- | --- | --- |
| Epitope | TC ♂ | TH ♂ | P-value | TC ♂ | TH ♂ | P-value |
| pY18 | 100.0 ± 10.2 | 141.3 ± 10.3 | P = 0.014 ° | 100.0 ± 5.6 | 79.0 ± 3.0 | P = 0.003 °° |
| pT181 | 100.0 ± 7.7 | 113.2 ± 6.3 | P = 0.203 | 100.0 ± 5.0 | 58.0 ± 4.3 | P = 0.001 °°° |
| pT214 | 100.0 ± 4.4 | 110.9 ± 6.5 | P = 0.211 | 100.0 ± 5.9 | 80.1 ± 5.1 | P = 0.021 ° |
| pS212/T214 | 100.0 ± 13.3 | 155.0 ± 20.0 | P = 0.050 ° | 100.0 ± 7.1 | 75.4 ± 11.0 | P = 0.089 |
| pS396 | 100.0 ± 5.2 | 127.7 ± 10.2 | P = 0.038 ° | 100.0 ± 7.4 | 68.9 ± 11.0 | P = 0.038 ° |
| pS404 | 100.0 ± 11.6 | 117.2 ± 15.6 | P = 0.415 | 100.0 ± 11.6 | 78.1 ± 8.4 | P = 0.143 |
| pS396/S404 | 100.0 ± 7.7 | 147.9 ± 11.2 | P = 0.005 °° | 100.0 ± 8.2 | 47.2 ± 9.5 | P = 0.001 °°° |
| pS422 | 100.0 ± 10.0 | 100.3 ± 6.7 | P = 0.980 | 100.0 ± 6.9 | 77.1 ± 1.6 | P = 0.004 °° |
| tau1 | 100.0 ± 6.0 | 98.0 ± 6.1 | P = 0.821 | 100.0 ± 8.2 | 91.4 ± 8.6 | P = 0.484 |
| Total tau | 100.0 ± 4.4 | 89.3 ± 4.7 | P = 0.128 | 100.0 ± 11.4 | 129.9 ± 5.7 | P = 0.028 ° |
|  | TC ♀ | TH ♀ | P-value | TC ♀ | TH ♀ | P-value |
| pY18 | 100.0 ± 7.9 | 104.3 ± 4.7 | P = 0.641 | 100.0 ± 8.3 | 141.0 ± 7.1 | P = 0.002 °° |
| pT181 | 100.0 ± 6.2 | 97.1 ± 2.7 | P = 0.661 | 100.0 ± 5.1 | 144.0 ± 13.1 | P = 0.014 °° |
| pT214 | 100.0 ± 5.8 | 105.9 ± 6.0 | P = 0.497 | 100.0 ± 5.4 | 131.9 ± 5.9 | P = 0.001 °°° |
| pS212/T214 | 100.0 ± 5.7 | 105.7 ± 9.4 | P = 0.621 | 100.0 ± 5.9 | 130.2 ± 8.4 | P = 0.011 ° |
| pS396 | 100.0 ± 7.6 | 82.2 ± 4.9 | P = 0.063 | 100.0 ± 8.4 | 128.6 ± 9.9 | P = 0.053 |
| pS404 | 100.0 ± 4.8 | 81.7 ± 8.8 | P = 0.116 | 100.0 ± 9.6 | 90.2 ± 7.5 | P = 0.428 |
| pS396/S404 | 100.0 ± 9.9 | 80.3 ± 2.8 | P = 0.062 | 100.0 ± 9.9 | 153.1 ± 8.6 | P = 0.001 °° |
| pS422 | 100.0 ± 10.1 | 79.3 ± 7.7 | P = 0.119 | 100.0 ± 6.8 | 167.4 ± 12.2 | P = 0.001 °°° |
| tau1 | 100.0 ± 4.2 | 95.8 ± 7.9 | P = 0.428 | 100.0 ± 8.3 | 86.6 ± 7.9 | P = 0.264 |
| Total tau | 100.0 ± 9.8 | 87.6 ± 4.5 | P = 0.254 | 100.0 ± 8.6 | 93.8 ± 6.9 | P = 0.584 |

### Sex-dependent effects of maternal HFD on hippocampal synaptic markers in adult THY-Tau22 offspring

To examine the effects of maternal HFD on synapses, we measured the protein level of presynaptic (SNAP25) and postsynaptic (NR2B) markers from hippocampal synaptosomal fractions of 7-month-old offspring. Immunoblots analysis revealed that maternal HFD significantly decreased NR2B (*P* < 0.01, Fig. 2A) and tended to decrease SNAP25 (*P* = 0.07, Fig. 2B) in THY-Tau22 male offspring, but not in female (Fig. 2C and 2D). Consistently, maternal HFD reduced the density of synaptophysin immunoreactivity in male offspring (*P* < 0.05, Fig. 3B), without affecting PSD95 immunoreactivity in both sexes (Fig. 3C, 3D and 3E). These findings suggest that maternal HFD induces hippocampal synaptic loss only in male offspring, pointing again to a sex-dependent effect.

**Figure 2.**
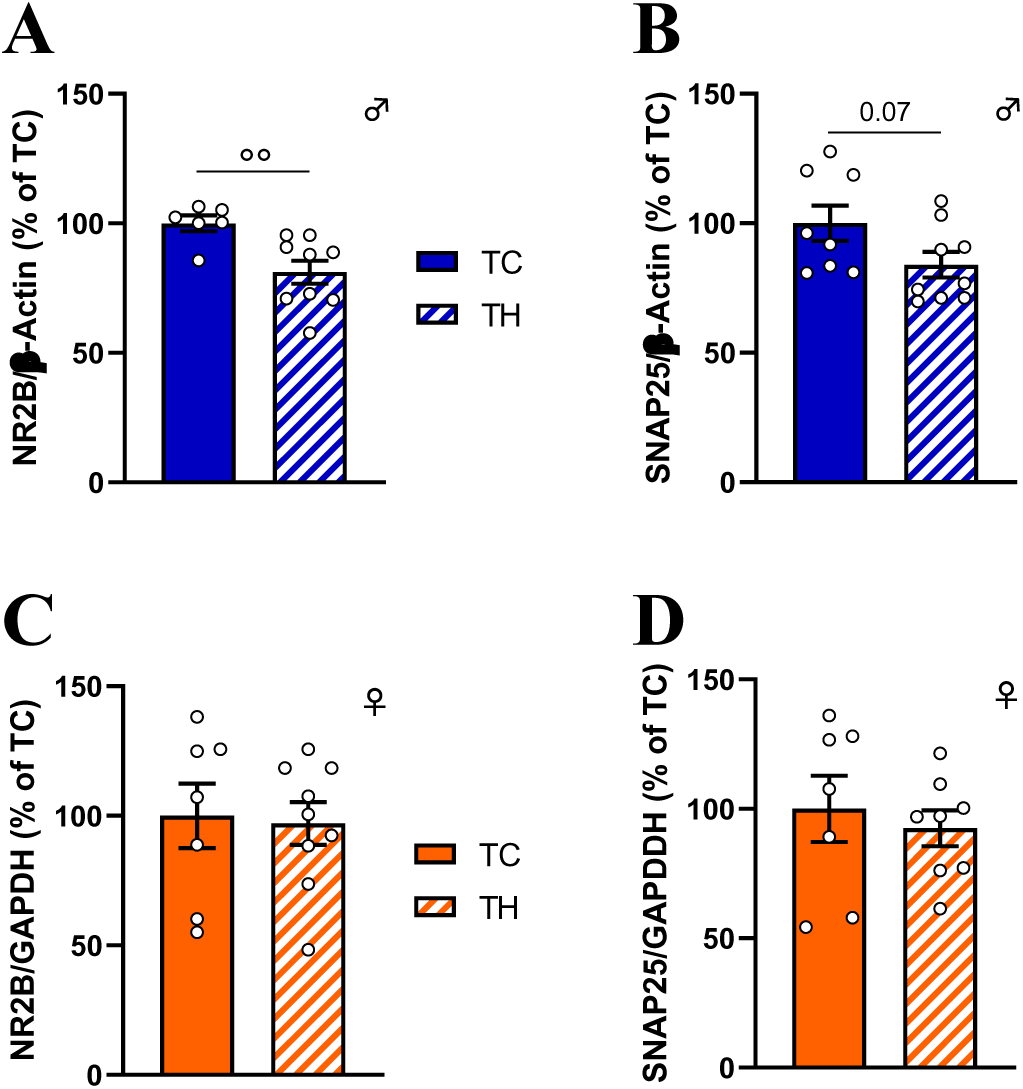
Effect of maternal HFD on hippocampal synaptic markers in adult THY-Tau22 offspring. Protein levels in fractions of hippocampal synaptosomes of 7-month-old TC and TH male and female mice were quantified using western blot. Quantification of the postsynaptic protein NR2B (**A** and **C**) and the presynaptic protein SNAP25 (**B** and **D**) in the hippocampal fractions of synaptosomes. °°*P* < 0.01, *P* = 0.07 versus TC mice using Student’s *t*-test. *n* = 6-9 per group. Values are expressed as mean ± SEM.

**Figure 3.**
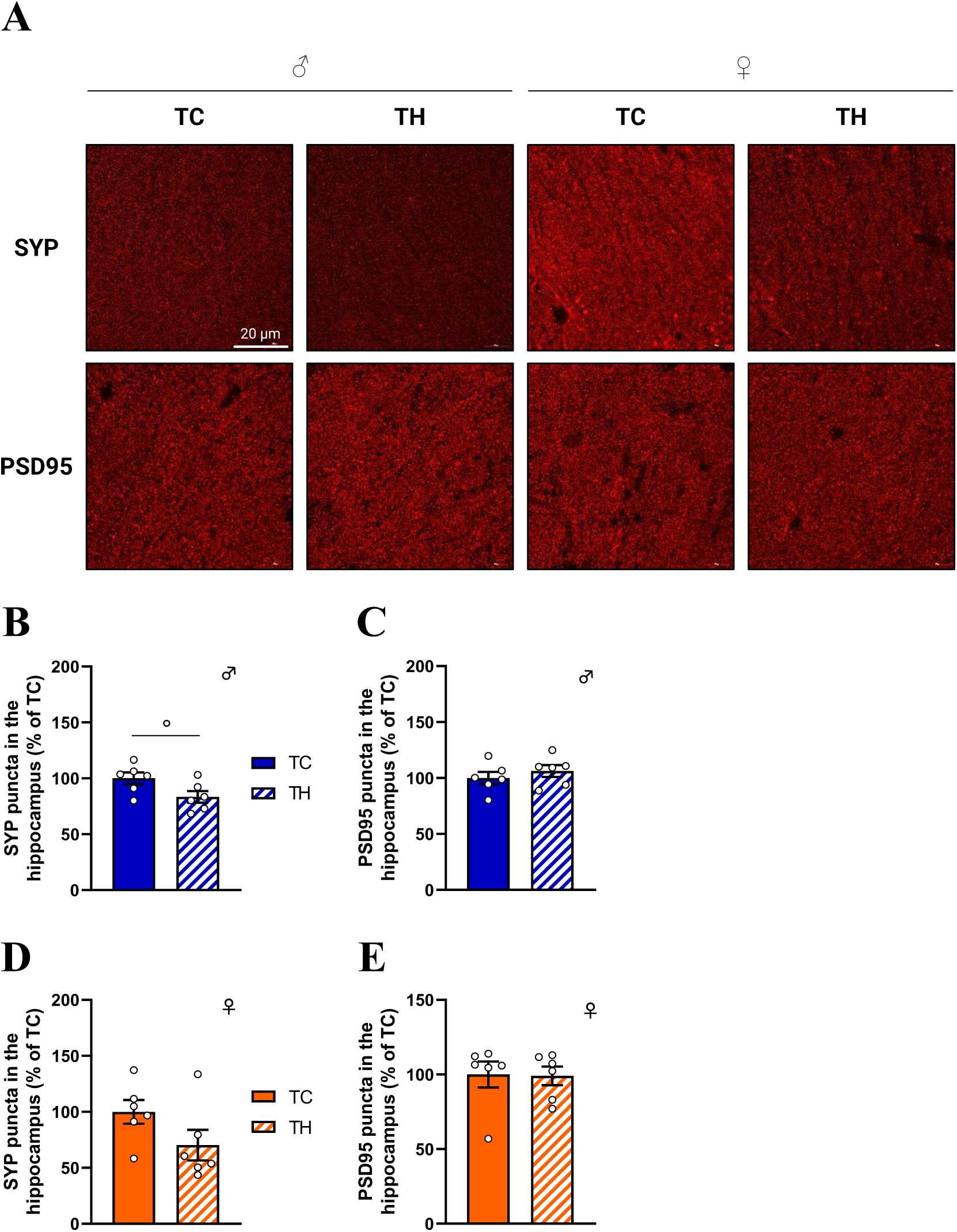
Effect of maternal HFD on hippocampal synaptic markers in adult THY-Tau22 offspring. (**A**) Characteristic immunofluorescence for presynaptic SYP and post-synaptic PSD95 markers (red) in the hippocampus of 7-month-old male and female TC and TH mice (**B** and **D**). Quantification of SYP (**C** and **E**) and PSD95 stainings. °*P* < 0.05 using a Student’s *t*-test. *n* = 6 per group. Values are expressed as mean ± SEM.

### Sex-dependent effects of maternal HFD on hippocampal neurogenesis in adult THY-Tau22 offspring

We investigated adult hippocampal neurogenesis (AHN) by injecting EDU 21 days before sacrifice at 7 months. Maternal HFD had no impact on the density of EDU-positive cells in the subgranular zone (SGZ) of the dentate gyrus of the hippocampus in male and female offspring (Fig. 4C and 4F). The differentiation of newly formed cells was analysed by co-labelling of EDU with a marker for immature (doublecortin, DCX) or mature (neuronal nuclear antigen, NeuN) neurons (Fig. 4A and 4B). Maternal HFD increased the proportion of NeuN-positive cells among EDU-positive cells only in female offspring (*P* < 0.05, Fig. 4G), this effect not being observed in male offspring (Fig. 4D). In addition, although not statistically significant, maternal HFD tended to reduce the proportion of EDU^+^/DCX^+^ immature neurons in female offspring (Fig. 4E and 4H). Finally, to finely study the impact of the diet on neuronal maturation, we performed a 3D reconstruction followed by a Scholl analysis of EDU^+^/DCX^+^ cells of the SGZ (Fig. 4I). Maternal HFD increased the distance between the soma and the furthest point reached by the dendritic arborization only in male offspring (*P* < 0.05, Fig. 4J and 4K). Altogether, these data reveal that maternal HFD modifies AHN in 7-month-old offspring in a sex-specific manner.

**Figure 4.**
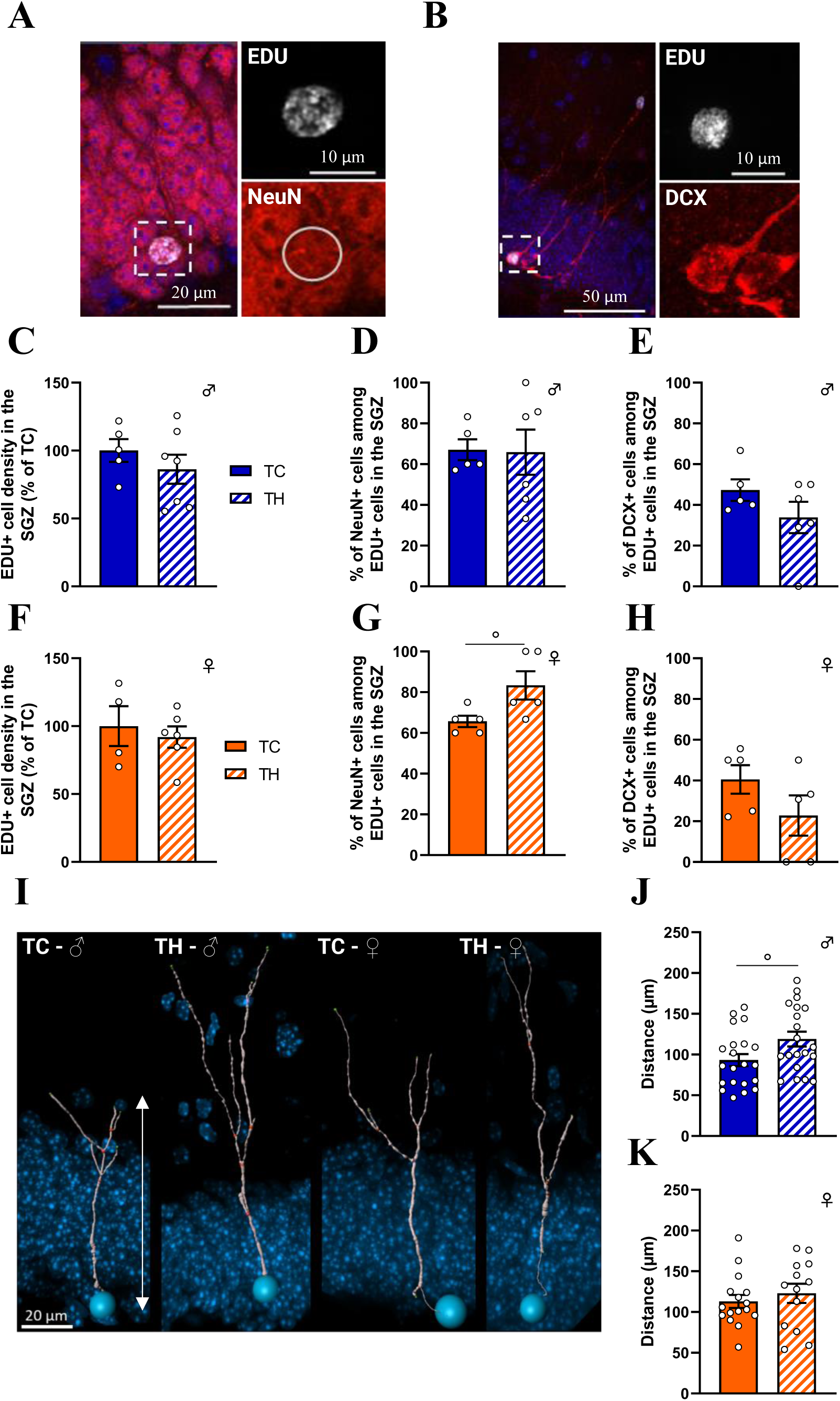
Effect of maternal HFD on adult hippocampal neurogenesis in adult THY-Tau22 offspring. Adult hippocampal neurogenesis (AHN) was analyzed by injecting a modified thymidine (EDU) three weeks before the sacrifice at 7 months of age. (**A**) Image showing an EDU^+^/NeuN^+^ cell (mature neuron marker, red) in the subgranular zone (SGZ). The circle surrounds the position of the EDU^+^ cell. DAPI (blue) represents cell nuclei. (**B**) Image showing an EDU^+^/DCX^+^ cell (immature neuron marker, red) in the SGZ. DAPI (blue) represents cell nuclei. (**C** and **F**) Quantification of the total number of EDU^+^ cells in the SGZ. (**D** and **G**) Quantification of the proportion of NeuN^+^ cells among EDU^+^ cells in the SGZ. (**E** and **H**) Quantification of the proportion of DCX^+^ cells among EDU^+^ cells in the SGZ. (**I**) Dendritic arborization of new neuronal cells (EDU^+^/DCX^+^ cells in the SGZ) was analyzed from confocal images using Imaris software. The 3D reconstruction from one representative cell per group is shown. Double arrow line represents the distance between the soma and the furthest point reached by the dendritic arborization. DAPI represented cell nuclei. (**J** and **K**) Quantification of this “distance”. °*P* < 0.05 versus TC mice using Student’s *t*-test. For **C**-**H** graphs, *n* = 6 mice per group. For **J** and **K**, each point represents one cell, i.e. *n* = 13-23 cells from six mice per group. Values are expressed as mean ± SEM.

### Sex-dependent effects of maternal HFD on the hippocampal transcriptome in adult THY-Tau22 offspring

Results of hippocampal RNA-seq indicated that maternal HFD tripled the number of significantly deregulated genes [DEGs; |log2 (fold-change)| > 0.26, adjusted *p*-value < 0.05] between 4 and 7 months of age in male offspring (1145 and 3264 deregulated genes, respectively under chow and HFD, Fig. 5A) and decreased it in female offspring (1413 and 280 deregulated genes, respectively under chow and HFD, Fig. 5B), again suggesting sex-related molecular differences. Thus, maternal HFD modifies the evolution of the hippocampal transcriptome over the age and the progression of tau pathology in a sex-dependent manner.

**Figure 5.**
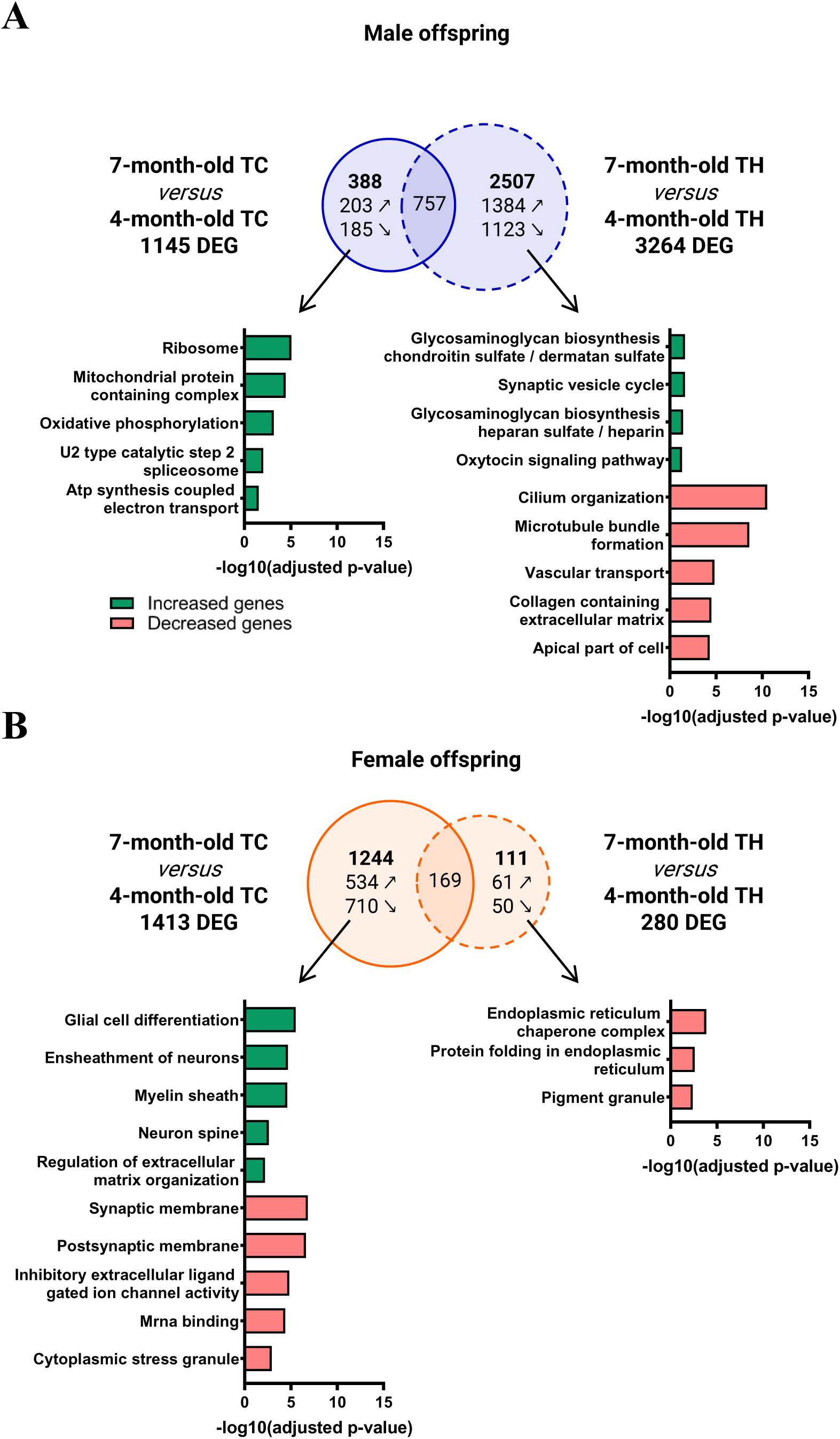
Effect of maternal HFD on hippocampal transcriptome in adult THY-Tau22 offspring. Hippocampal transcriptome analysis was performed using RNA sequencing in male (**A**) and female (**B**) THY-Tau22 offspring aged of 4 and 7 months of age. Venn diagrams indicate the number of significantly deregulated genes [|log2 (fold-change)| > 0.26, adjusted p-value < 0.05] exclusively between 4- and 7-month-old animals on chow and HFD during lactation, including the overlap of deregulated genes across the two comparisons. Histograms show the results of over-representation analysis (ORA) of the deregulated genes, categorized by Kyoto encyclopedia of genes and genomes (KEGG) and gene ontology (GO) biological processes (BP), cellular components (CC) and molecular functions (MF). ORA was performed separately on genes that were exclusively deregulated between 4 and 7 months under chow, with genes showing increased expression indicated in green and those with decreased expression showed in red (two separate analyses). The same analysis was then conducted for genes exclusively deregulated under HFD. The top five enriched pathways are shown.

Then, we performed an Over Representation Analysis (ORA) to identify the pathways enriched with genes showing an increased or decreased expression exclusively by the chow diet (age effect) or HFD (age x diet effect). In male offspring (Fig. 5A), while no significant enriched pathways was observed when analysing the 185 genes exclusively decreased by chow diet, the 203 genes increased were mainly associated with ribosomes and mitochondrial respiration pathways. The 1384 genes exclusively increased by maternal HFD were enriched in pathways related to glycosaminoglycan biosynthesis pathways and synaptic vesicle cycle, while the 1123 decreased were associated with cilium, vascular transport and extracellular matrix pathways (Fig. 5A). The ORA performed in female offspring revealed a profound sex-related effect. Indeed, the 534 genes exclusively increased by aging were enriched in pathways related to glial cell differentiation, myelination and extracellular matrix, while the 710 decreased were associated with synaptic plasticity. Notably, while there was no significant enriched pathways when analysing the 61 genes exclusively increased by maternal HFD, the 50 genes decreased were associated with endoplasmic reticulum pathways (Fig. 5B).

### Sex-dependent effects of maternal HFD on the hippocampal proteome in adult THY-Tau22 offspring

Mass spectrometry analyses, performed in the same animals that have been examined by RNA-seq, revealed that very few of the genes deregulated between 4 and 7 months of age were also affected at the protein level, irrespective of diet or sex (Supplementary Fig. 7). Hippocampal proteome data analysis, as for the transcriptome, showed an increase of the number of significantly deregulated proteins in male offspring between 4 and 7 months, with 75 and 229 deregulated proteins (including 53 shared), respectively under chow diet and HFD (Fig. 6A). In contrast to what was observed in the transcriptome study, the diet slightly increased the number of deregulated proteins in female offspring between 4 and 7 months, with 9 proteins showing a modified level under chow diet and 29 under HFD including 5 in common (Fig. 6B). Furthermore, the proteome evolution with age was more impacted in male offspring, particularly under maternal HFD (Fig. 6).

**Figure 6.**
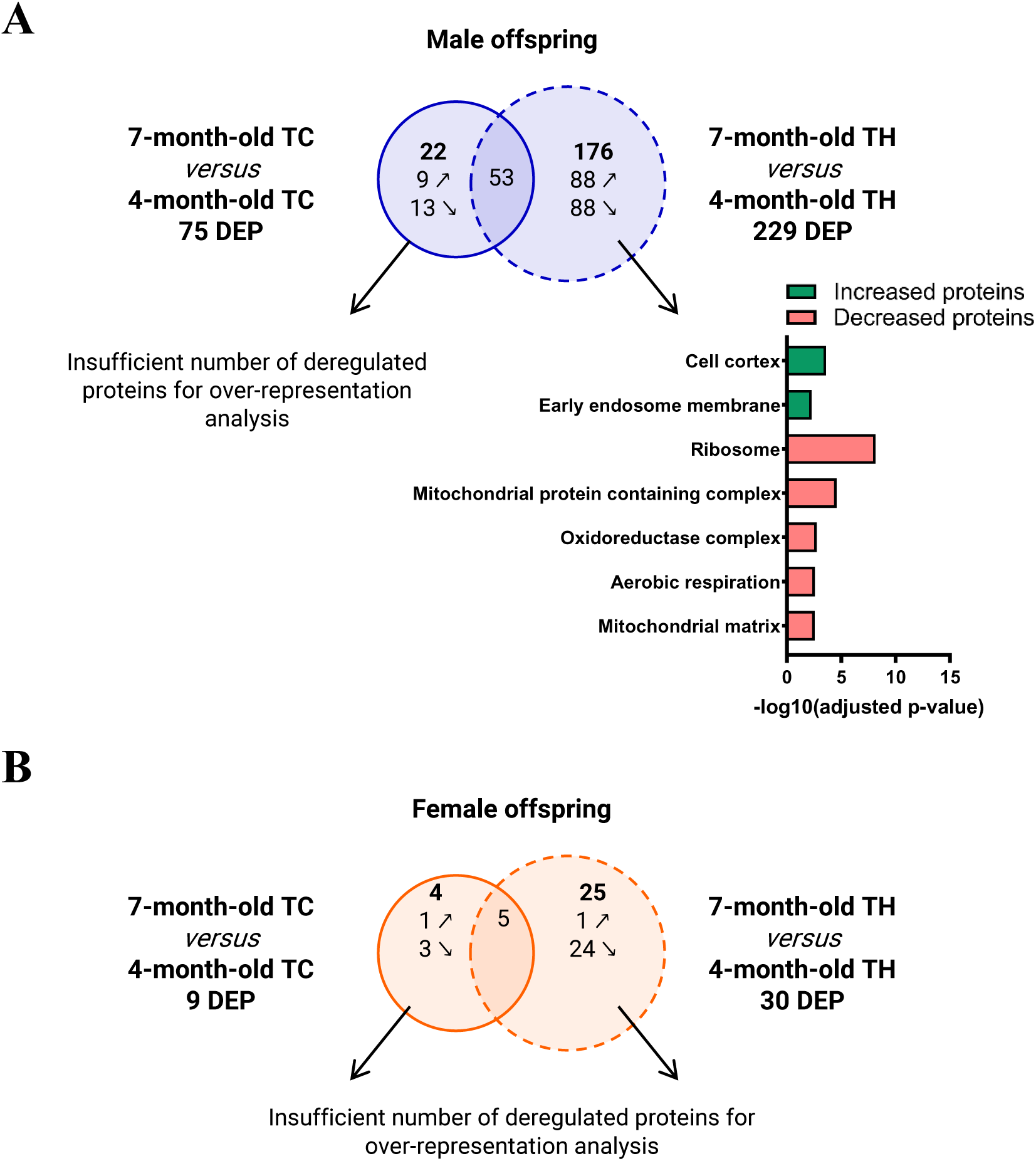
Effect of maternal HFD on hippocampal proteome in adult THY-Tau22 offspring. Hippocampal proteome analysis was performed using mass spectrometry in male (**A**) and female (**B**) THY-Tau22 offspring aged at 4 and 7 months of age. Venn diagrams indicate the number of significantly deregulated proteins [|log2 (fold-change)| > 0.26, adjusted p-value < 0.05] exclusively between 4- and 7-month-old animals on chow and HFD during lactation, including the overlap of deregulated proteins across the two comparisons. Histograms show the results of over-representation analysis (ORA) of the deregulated genes, categorized by Kyoto encyclopedia of genes and genomes (KEGG) and gene ontology (GO) biological processes (BP), cellular components (CC) and molecular functions (MF). ORA was performed separately on genes that were exclusively deregulated between 4 and 7 months under chow, with proteins showing increased expression indicated in green and those with decreased expression showed in red (two separate analyses). The same analysis was then conducted for proteins exclusively deregulated under HFD. The top five enriched pathways are shown. When fewer than 25 proteins were deregulated, ORA was not performed, and manual annotation was conducted instead (Supplementary Tables 5, 6 and 7).

ORA showed that in male offspring, 7 of the 13 proteins exclusively decreased by the chow diet with age were associated with the synapse (Supplementary Table 5). Furthermore, the 88 proteins increased exclusively by maternal HFD were linked to the cortex cell and endosomes, while the 88 decreased were enriched in mitochondrial and ribosomal pathways (Fig. 6A). In female offspring, although there were not enough proteins to perform an ORA, manual annotations indicated that half of the proteins exclusively decreased by maternal HFD were related to mitochondria or synapse (Supplementary Tables 6 and 7).

### Sex-dependent effects of maternal HFD on hippocampal regulons in adult THY-Tau22 offspring

Finally, in order to get an overview and to integrate hippocampal transcriptomic and proteomic data, we studied the regulome of 4- and 7-month-old offspring. At 4 months, while only one regulon was found to be deregulated by maternal HFD in female offspring (Bclaf1, FDR < 0.01), seven regulons were significantly downregulated in males (Trim28, Sfpq, Nono, Mecp2, Ctnnb1, Ctbp1, Bcalf1), suggesting an earlier impact in males (Fig. 7A). All these regulons were found associated with decreased expression by maternal HFD. In contrast, at 7 months, five regulons were found modified by maternal HFD in both males and females (Prdm16, Mecp2, Ctnnb1, Ctbp1, Bclaf1), two only in males (Trim28 and Smarca4) and two only in females (Sfpq and Nono). Except Trim28 and Smarca4, these regulons exhibit decreased expression of their genes by maternal HFD (NES < 0, Fig. 7A). Further, enrichment analysis indicated that the genes encompassed within the regulons deregulated by maternal HFD at 4 months in males and 7 months in both sexes were linked to the mitochondrial, ribosomal, gliogenesis and myelination pathways (Fig. 7B). Interestingly, genes associated with Trim28 and Smarca4 regulons, found upregulated only in males, were associated with immune response, suggesting that this function is altered differently according to sex.

**Figure 7.**
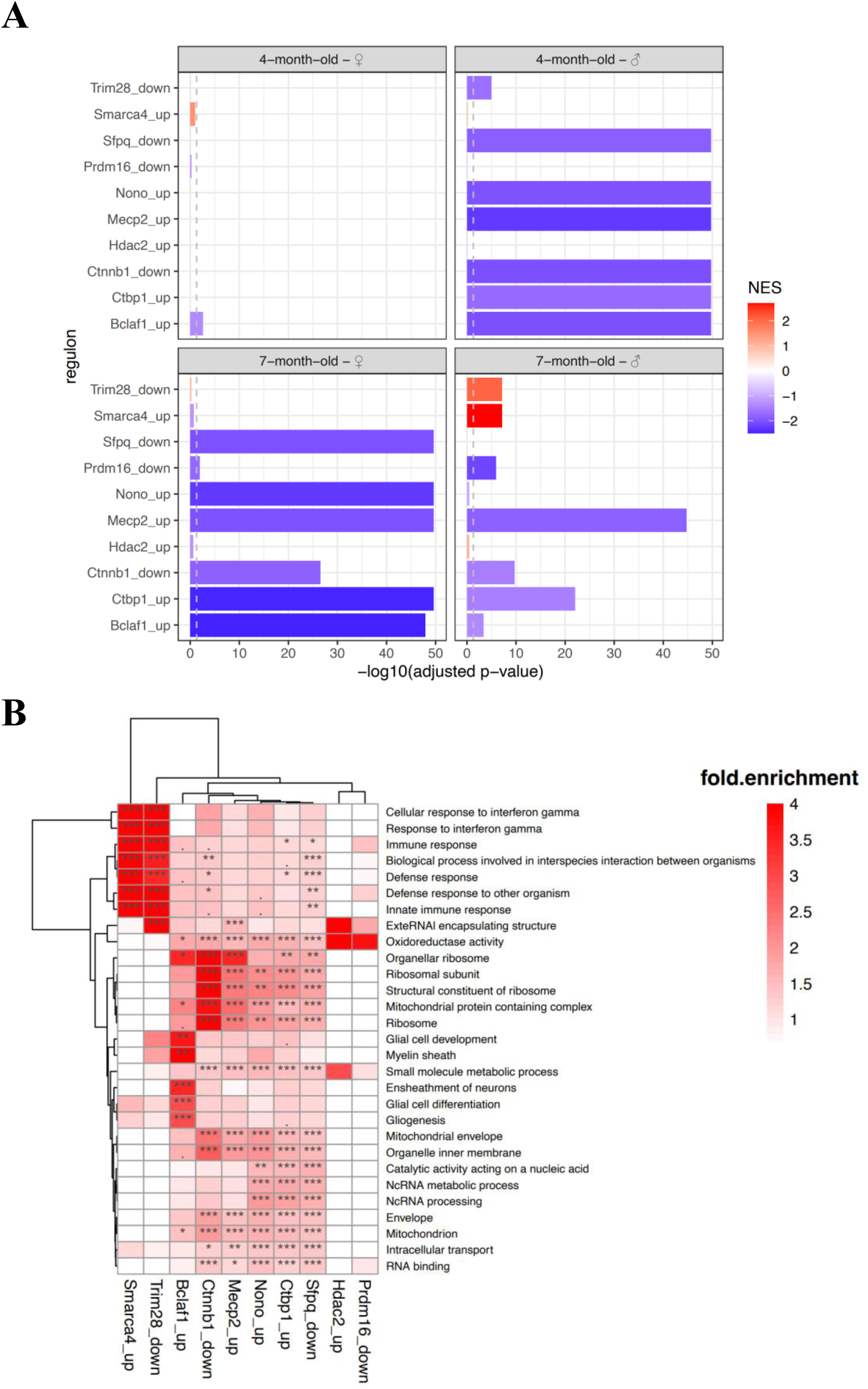
Identification of deregulated regulons by maternal high-fat diet in adult THY-Tau22 offspring. Regulons were identified using hippocampal transcriptome and proteome data from 4- and 7-month-old THY-Tau22 animals. (**A**) Bar plots show regulons that are significantly enriched (adjusted p-value < 0.01) in 4- and 7-month-old male and female animals that have received maternal HFD during lactation. A positive NES means that the regulon is enriched in upregulated genes. In the regulon name, “up” and “down” refer to the activating or inhibiting effect of the transcription factor on the target genes, respectively. (**B**) Over-representation analysis was conducted for each identified regulon. The heatmap displays the top five gene ontology (GO) terms, including biological processes, molecular functions, cell components, as well as Kyoto encyclopedia of genes and genomes (KEGG) pathways..*P* < 0.1, \**P* < 0.05, \*\**P* < 0.01, \*\*\**P* < 0.001.

## Discussion

Many epidemiological and experimental studies show that perturbations of the perinatal environment can contribute to the development of diseases in adult offspring.^33,54,69^ In particular, recent studies reported that maternal malnutrition is associated with increased risk of metabolic and cognitive disorders in the offspring,^38,40,70,71^ with long-term alterations in the hippocampus, a brain structure involved both in cognition but also the regulation of metabolism.^41,44,46,72,73^. Since metabolic disorders are risk factors for neurodegenerative diseases such as AD,^9,12^ it is conceivable that maternal malnutrition may contribute to its development. However, only few and contradictory studies have investigated the putative links between maternal malnutrition and AD in offspring,^56–58^, and none of them have addressed sex-dimorphism. In the present study, we show, using the THY-Tau22 mice, a model of AD-like tauopathy, that maternal HFD applied only during lactation worsens tau pathology, synaptic loss and memory deficits with long-term consequences on gene and proteins expression in the offspring. Interestingly, the effects observed exhibit sex dependency with male offspring being more impacted than females.

At the metabolic level, we show that maternal HFD increases body weight in both male and female THY-Tau22 offspring at weaning, suggesting enhanced lactation efficiency. This effect, consistent with a previous meta-analysis in WT offspring,^74^ was also observed in littermate controls,^44^ indicating that it results primarily from maternal HFD. Studies suggest that the weight gain could be due to changes in milk composition,^75–78^ as well as modification in maternal behaviour.^79–81^ In adulthood, body weight was normalized indicating a transient effect of maternal HFD. This result is surprising because maternal HFD exposure that promotes increased body weight gain during lactation is usually associated with metabolic disorders and obesity in the adulthood.^71,82–84^ Although we did not observe major disturbances in plasma level of free fatty acid, cholesterol, triglyceride, or insulin, maternal HFD leads to mild glucose intolerance in 4-month-old male offspring, as previously shown in control offspring and other studies on sex dimorphism,^44,85^ suggesting that young adult males are more vulnerable to the maternal diet.

THY-Tau22 mice develop spatial memory impairments starting at 7-8 months of age.^59,60^ Here, we show that maternal HFD accelerates the onset of cognitive impairments as early as 6 months of age in both sexes. Using the Y-maze, we demonstrated a short-term spatial memory alteration in both male and female offspring following maternal HFD. Additionally, using the Barnes maze, we showed that maternal HFD also affects long-term reference spatial memory in male offspring, with no impact on females. To our knowledge, this sex-specific effect has never been reported in this context, although it aligns with previous studies reporting memory impairment in male wild-type animals exposed to maternal HFD.^44,86,87^ Our data are consistent with findings in humans as well as preclinical data from the 3xTgAD model, which revealed that maternal obesity and/or malnutrition during the perinatal period exacerbates memory deficits in offspring.^38–40,56,57^ However, using the same 3xTgAD model, a beneficial effect was also reported when the diet was applied only during gestation,^58^ suggesting that the lactation period, which corresponds to the peak of brain development in rodent,^32^ is particularly sensitive to the deleterious effects of the maternal diet.

Interestingly, diet-induced memory impairment is associated with an increase in hippocampal tau pathology as early as 4 months of age in male offspring, while this is observed later, at 7 months, in female offspring. This temporal shift could explain the absence of long-term memory deficit in females, consistent with an increased vulnerability to the diet in males and the instrumental role of tau pathology in cognition.^88^ It is possible that females are protected by estrogens which have been suggested to exert a neuroprotective role against tau pathology.^89–93^ The increase in tau pathology induced by maternal HFD does not seem to be linked to major neuroinflammatory changes either. However, *Trem2* expression, which plays a crucial role in AD pathogenesis,^94^ is slightly increased by maternal HFD at 4 months in male offspring, and two maternal HFD-increased regulons (Trim28 and Smarca4) are associated with the immune response, highlighting a stronger effect in males. This would be linked to tau pathology development considering the vicious circle between neuroinflammatory processes and tau pathology development.^68^

We also investigated synaptic markers and AHN - two processes sensitive to tau pathology^95–98^ and maternal malnutrition,^42,45–47^ both of presumably explaining memory deficits induced by maternal HFD in tau offspring. In the synaptosomal fractions of the hippocampus from male TH offspring, we observed a decrease in the protein levels of SNAP25 and NR2B, involved in presynaptic exocytosis and glutamatergic signalling, respectively.^99,100^ In the same mice, we also detected a presynaptic alteration thanks to synaptophysin staining. These alterations may be linked to the toxic effects of tau pathology on synapses, as well as to disruptions in the extracellular matrix and cytoskeleton, as highlighted by omics data.^101–104^ The absence of changes in the expression of these synaptic markers in female TH offspring may reflect a compensatory mechanism, which could partly explain the lack of long-term memory impairment in these mice. Moreover, it has been shown that synaptic alterations can affect neuronal maturation,^105,106^ suggesting that synaptic loss may also contribute to the AHN modifications observed in male TH offspring. Indeed, the dendrites of newly generated neurons in these animals extend deeper into the molecular layer as compared to control mice, suggesting a remodelling of dendritic tree. Since increased dendritic arborisation has been associated with beneficial effects on memory in several studies,^107,108^ this remodelling may represent a compensatory mechanism, although it may be insufficient to counteract the effects of maternal HFD exposure given the impaired memory observed. Alternatively, a potential deleterious effect cannot be excluded. Indeed, a previous study performed in an amyloid mouse model reported an initial acceleration of new neuron growth, followed by a long-term slowdown compared to controls.^106^ Furthermore, modifications in AHN are also observed in female TH offspring. Specifically, they exhibit a higher proportion of mature neurons in the subgranular layer of the hippocampus. In line, omics data reveal a deregulation of genes related to cell differentiation only in females. Similar to male offspring, this increase may reflect a compensatory mechanism aimed at mitigating cognitive decline. Further studies are needed to decipher whether the mechanisms involved are beneficial or detrimental. Collectively, these data reinforce the idea of a sex-dependent effects of maternal HFD, with a greater or earlier vulnerability in male offspring.

Finally, we performed a time-course investigation at both 4-(at the beginning of tau pathology) and 7-month-old (just before onset of cognitive disorders) male and female THY-Tau22 mice, using multi-omics analysis combining transcriptome, proteome and regulome analysis, to identify the biological pathways affected by maternal HFD in adult offspring. To our knowledge, this type of comprehensive and unbiased approach has never been conducted in AD and tauopathy mouse models in these experimental settings. We observed that maternal HFD increases the number of significantly dysregulated genes during the progression of tau pathology in males, but decreases it in females. This reduction may reflect a compensatory mechanism to mitigate the detrimental effects of the diet or alternatively may be deleterious itself. In contrast, maternal HFD increases the number of deregulated proteins in both males and females, with a greater number of deregulated proteins in males, highlighting a more pronounced impact of maternal HFD. Similar to what has been described in a multi-omics analysis of AD brain patients,^109^ the deregulated genes and proteins differ considerably, emphasizing the importance of investigating both the transcriptome and the proteome. This observation aligns with the changes in genes and proteins associated with ribosomes in male offspring, suggesting an alteration in translation. In addition to the biological pathways previously detailed, our data reveal that maternal HFD deregulates genes and proteins associated with the cilium and mitochondria, specifically in male offspring. Indeed, we found a reduction in cilium-associated genes in male TH offspring between 4 and 7 months. The primary cilium, a microtubule-based sensory organelle found on most cell types, including neurons and astrocytes, has been suggested to be a mediator of neural stem cells activity, neurogenesis, neuronal maturation and maintenance, as well as to play a role in AD development.^110–112^ Thus, alterations in microtubules could impair primary cilia, and lead to long-term memory deficits observed in male TH mice, as suggested by previous works.^113,114^ Interestingly, genes associated to cilium pathways were also found deregulated in their WH littermate controls, which also develop memory alterations.^44^ Moreover, proteomic data revealed a decrease of mitochondria-associated proteins between 4 and 7 months, specifically in male TH mice. These proteins are involved in oxidative phosphorylation, Krebs cycle, ketone metabolism and beta-oxidation, indicating impaired energy metabolism and reduced ATP production. Mitochondrial dysfunction, linked to cognitive decline, has also been observed in tauopathy models where tau pathology impairs mitochondrial axonal transport, thereby altering synaptic plasticity due to energy deficiency.^115–117^. In line, maternal stress models also showed increased amyloid pathology, memory deficits, and altered synaptic mitochondrial proteins, pointing out to mitochondria as a key target of perinatal disturbances in AD models.^118^ It is noteworthy that the effect of the diet on mitochondrial proteins is also present in female offspring, although it is weaker and involves different proteins. Finally, we showed that the regulons modified by maternal HFD, and which target mitochondria and ribosomes, are similar in both sexes but are delayed in females. These results suggest that the distinct effects of maternal HFD observed in male and female offspring are indeed more related to a temporal shift rather than distinct sex-specific alterations.

In conclusion, the present data support the view that maternal malnutrition is involved in the development of tauopathy and associated cognitive decline in a sex specific manner, male offspring being earlier impacted than females. In line with other studies,^56,57^ our work shows that the perinatal environment may participate to the programming of AD and tauopathies and that a healthy food intake during the perinatal period may protect from the development of neurodegenerative diseases.

## Acknowledgements

We thank the animal core facility (animal facilities of University of Lille) of “Plateformes lilloises en Biologie Santé” (PLBS, US 41 – UAR 2014), and more particularly the Specific Pathogen Free platform. We thank the In vivo Imaging and Functional Exploration platform (LiiFE, PLBS, US 41 – UAR 2014) where behavioural assessment was performed. We thank Myriem Tardivel, Antonino Bongiovanni and Sarah Gabut for their help on the Zeiss confocal microscope and Imaging analysis from the Photonic Microscopy Core BioImaging Center (BiCel, PLBS, US 41 – UAR 2014). We also thank the Bioinformatic, Bioanalysis and Biostastistic plateform (BiLille, PLBS, US 41 – UAR 2014). Sequencing was performed by the GenomeEast platform, a member of the “France Genomique” consortium (ANR-10-INBS-0009). We thank the OrganOmics platform of PRISM Inserm U1192 which is recognized and supported by the University of Lille and, the Infrastructure PROFI (https://www.profiproteomics.fr/) and the GIS IbiSA (https://www.ibisa.net/). The OrganOmics platform (Villeneuve d’Ascq, France) is also supported by Region Hauts de France and FEDER funding.

## Supplementary material

Supplementary material is available at *Brain* online.

## Data availability

Sequencing data have been deposited in the NCBI’s Gene Expression Omnibus (GEO) database (GSE297502). Mass spectrometry proteomics data have been deposited to the ProteomeXchange Consortium via the PRIDE partner repository with the database identifier PXD063974.^119^ Regulomic data are available on https://github.com/AlexandrePelletier/Perinatau/tree/main. The other data that support the findings of this study are available on request to corresponding authors.

## Supplementary figures, tables and legends

**Supplementary Figure 1.**
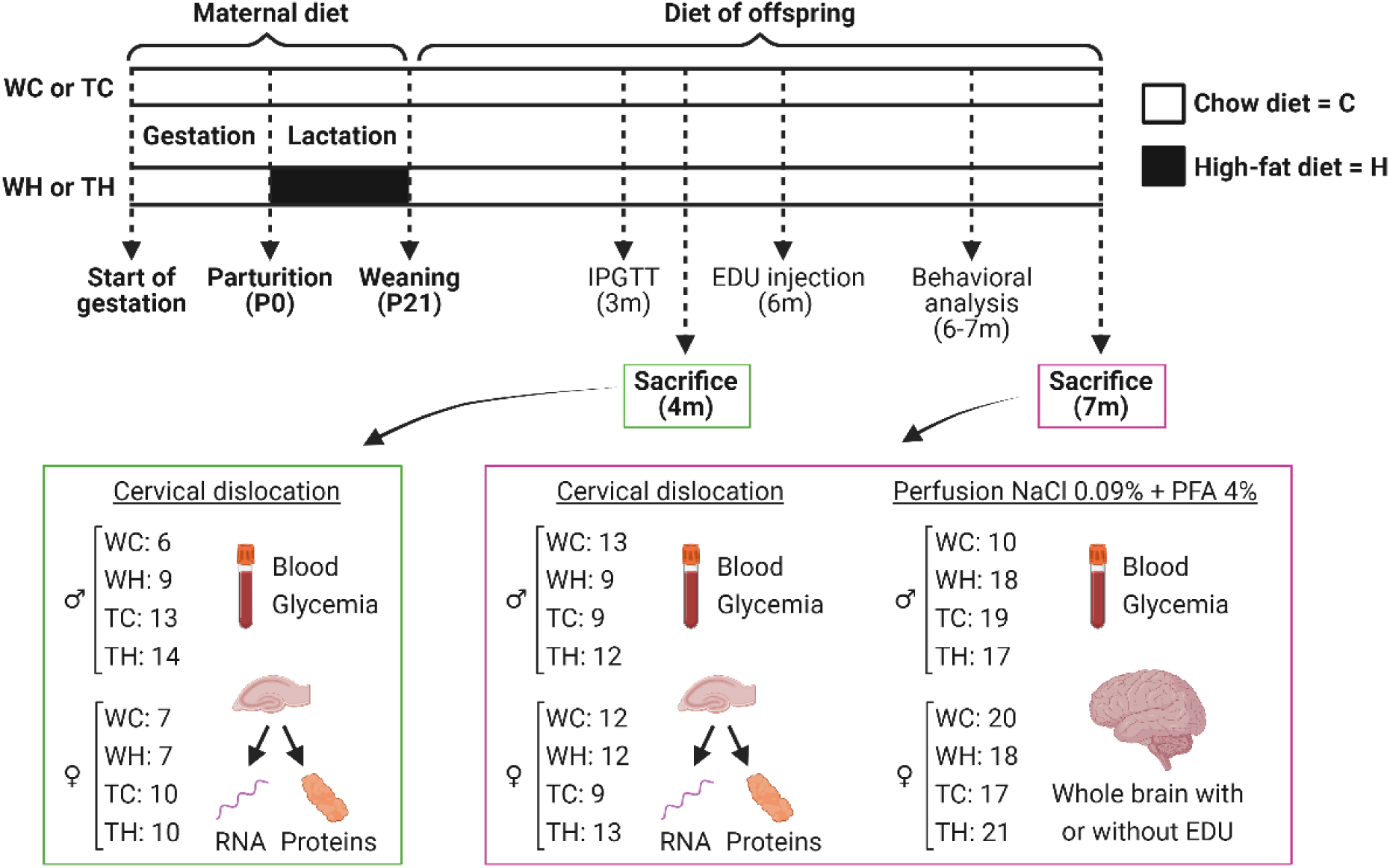
Experimental protocol. C57Bl6/J dams were fed a chow diet (13.5% fat, C; *n* = 28 dams) or a high-fat diet (HFD; 58% fat, H; *n* = 28 dams) during lactation, generating four offspring groups: THY-Tau22 Chow (TC), THY-Tau22 (TC) and their littermate WT Chow (WC) and WT (HFD) littermate controls. At weaning (postnatal day 21, P21), offspring returned to a standard diet (8.4% fat) until sacrifice at 4 or 7 months of age. An intraperitoneal glucose tolerance test (IPGTT) was performed in 3-month-old animals. Injection of 5-ethynyl-2’-deoxyuridin (EDU) and behavioral tests were performed at 6 and 6-7 months of age, respectively. The number of individuals per group, the method of sacrifice and the biological material recovered are indicated under the experimental protocol. Figure created with BioRender.com.

**Supplementary Figure 2.**
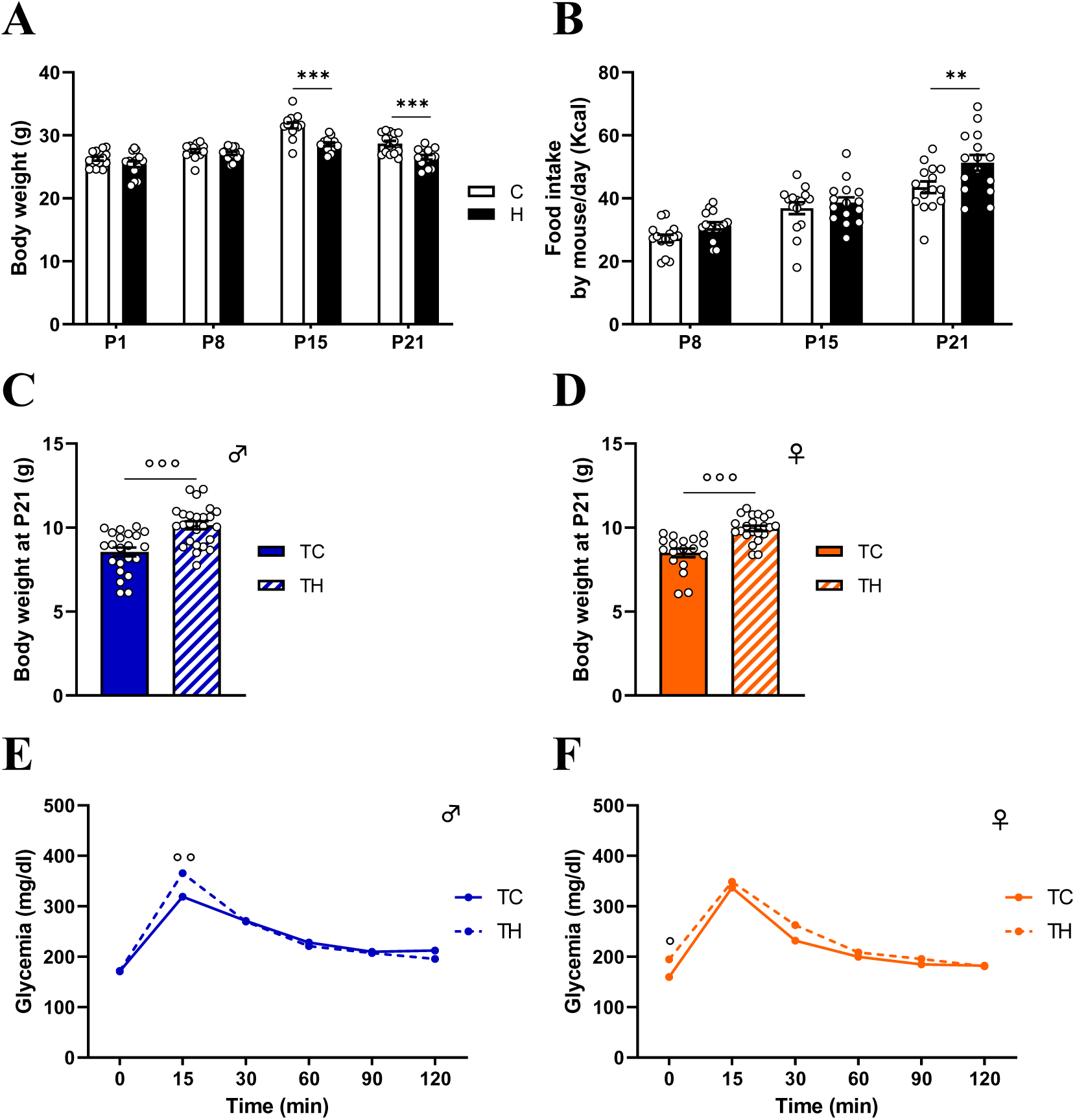
Effect of maternal HFD on metabolic parameters in dams and THY-Tau22 offspring. (**A**) Body weight of dams fed with Chow (C) or High-Fat (H) diets during lactation at different postnatal days (P1, P8, P15 and P21). (**B**) Food intake of dams fed with C or H diets during lactation at P8, P15 and P21. (**C** and **D**) Body weight of males (C) and female (D) offspring at weaning (P21). (**E** and **F**) Intraperitoneal glucose tolerance test was performed in 3-month-old in male (E) and female (F) offspring. The graphs show the evolution glycaemia at different time points (0, 15, 30, 60, 90 and 120 minutes following glucose injection) in male and female offspring. \*\**P* < 0.01, \*\*\**P* < 0.001, °*P* < 0.05, °°*P* < 0.01, °°°*P* < 0.001 using two-way ANOVA followed by Sidak’s *post hoc* test (**A**, **B**, **E** and **F**) or Student’s *t*-test (**C** and **D**). *n* = 14-19 per group. Values are expressed as mean ± SEM.

**Supplementary Figure 3.**
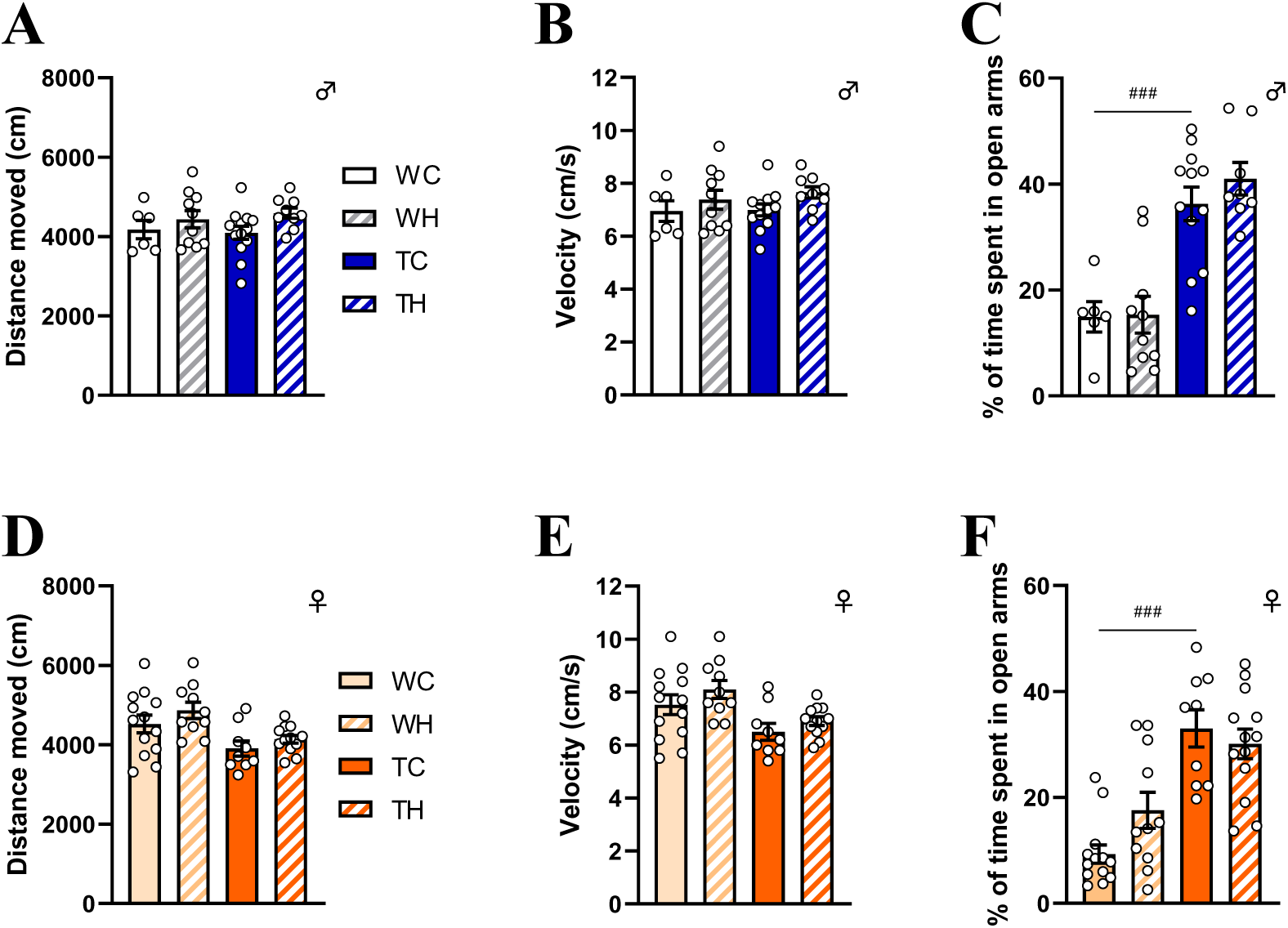
Effect of maternal HFD on spontaneous activity and anxiety-like behavior in 7-month-old THY-Tau22 offspring. (**A** and **D**) Distance moved and (**B** and **E**) velocity observed using actimetry to study spontaneous activity. (**C** and **F**) Percentage of time spent in open arms in the elevated plus maze test to study anxiety-like behavior. ##*P* < 0.01, ###*P* < 0.001 versus WC mice using one-way ANOVA followed by Tukey’s *post hoc* test. *n* = 6-14 per group. Values are expressed as mean ± SEM.

**Supplementary Figure 4.**
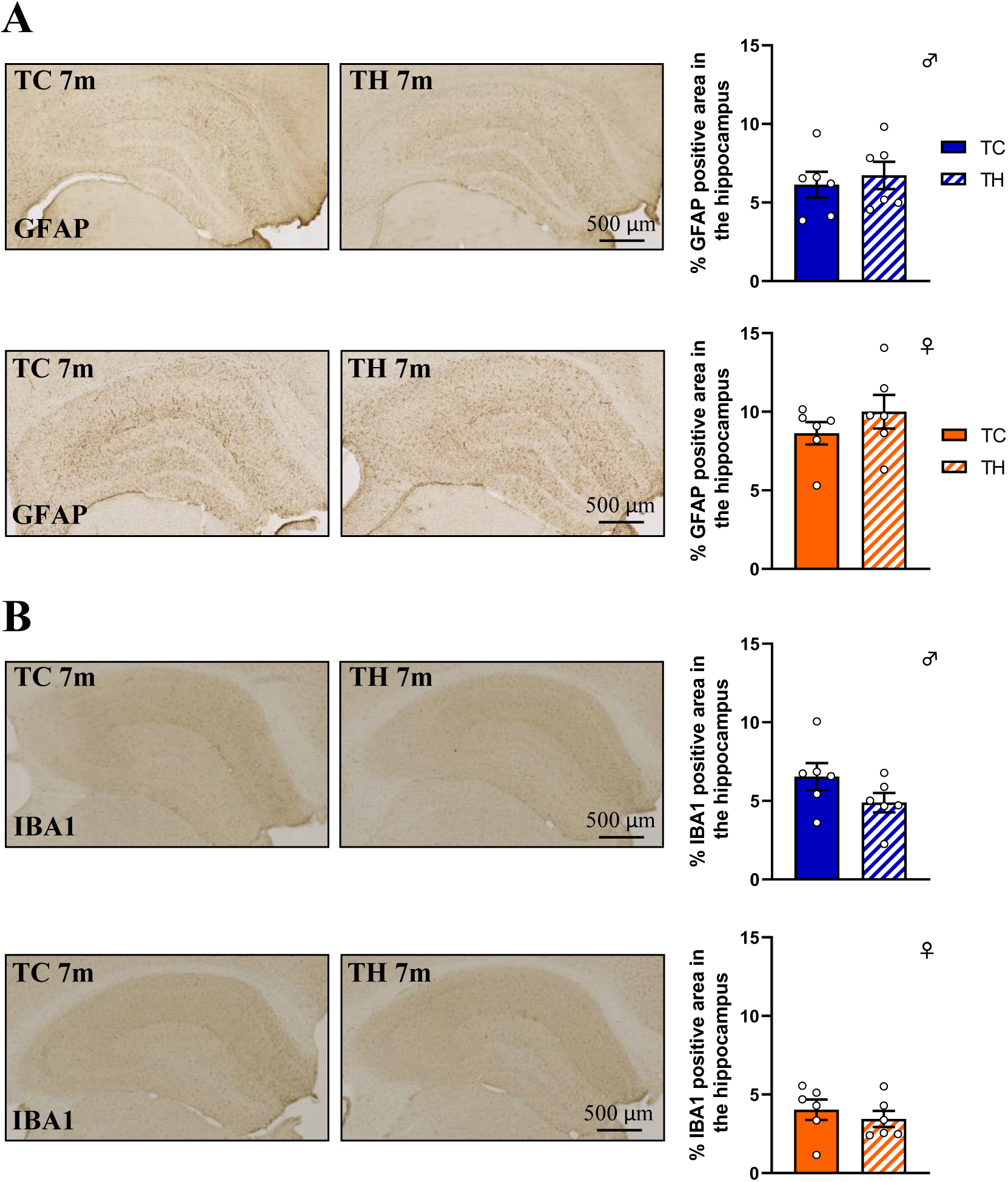
Effect of maternal HFD on hippocampal astrocytes and microglia in 7-month-old THY-Tau22 offspring. (**A**) Representative images of anti-GFAP (astrocytic marker) and (**B**) anti-IBA1 (microglial marker) immunohistochemistries and related quantifications of the staining area in the hippocampus. *n* = 6 per group. Values are expressed as mean ± SEM.

**Supplementary Figure 5.**
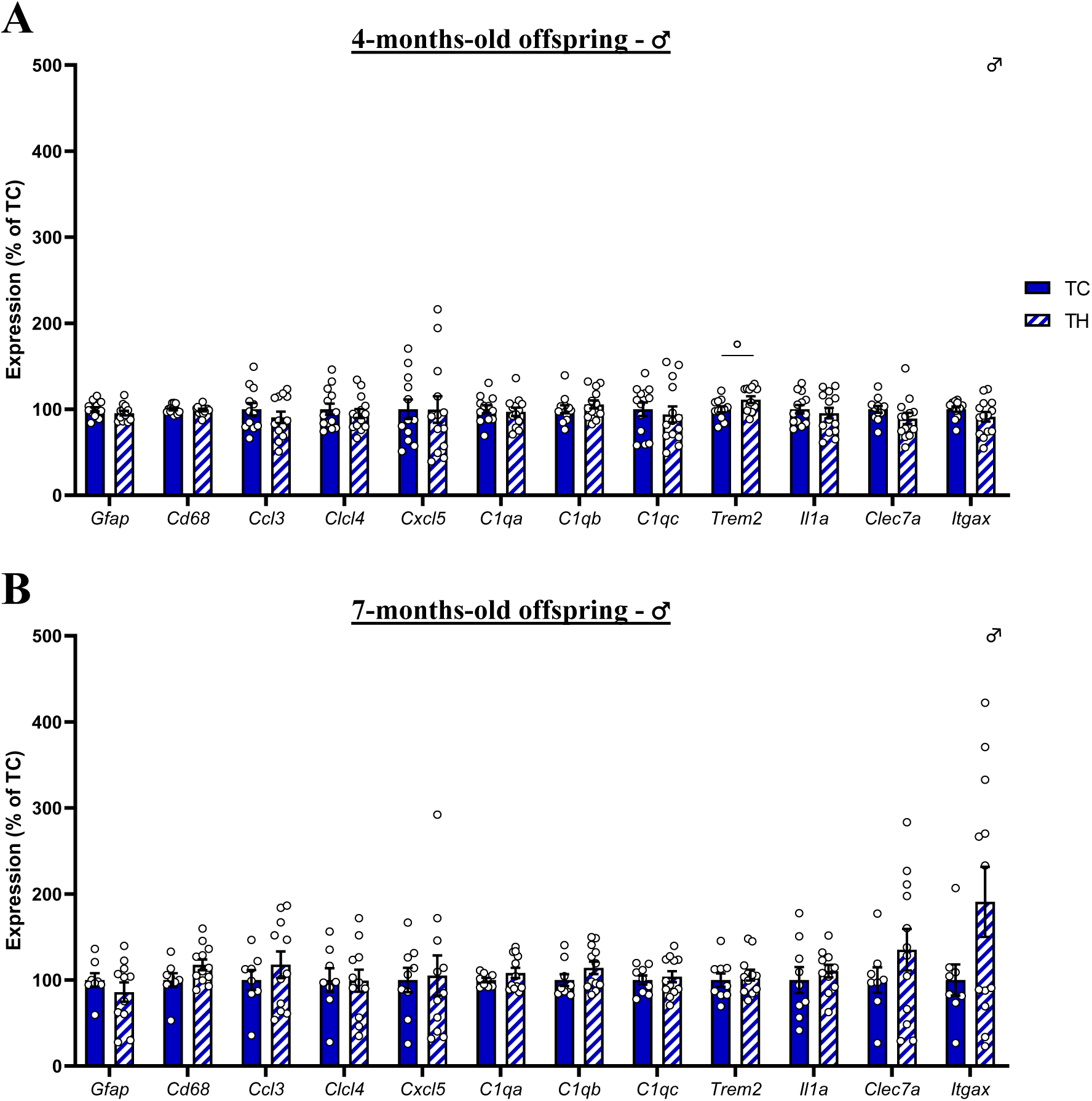
Effect of maternal HFD on hippocampal neuroinflammation in adult male THY-Tau22 offspring. (**A** and **B**) Expression of neuroinflammatory genes was measured by quantitative PCR in the hippocampus of 4-month-old and 7-month-old maleTHY-Tau22 offspring. °*P* < 0.05 versus TC mice using Student’s *t*-test. *n* = 6-14 per group. Values are expressed as mean ± SEM.

**Supplementary Figure 6.**
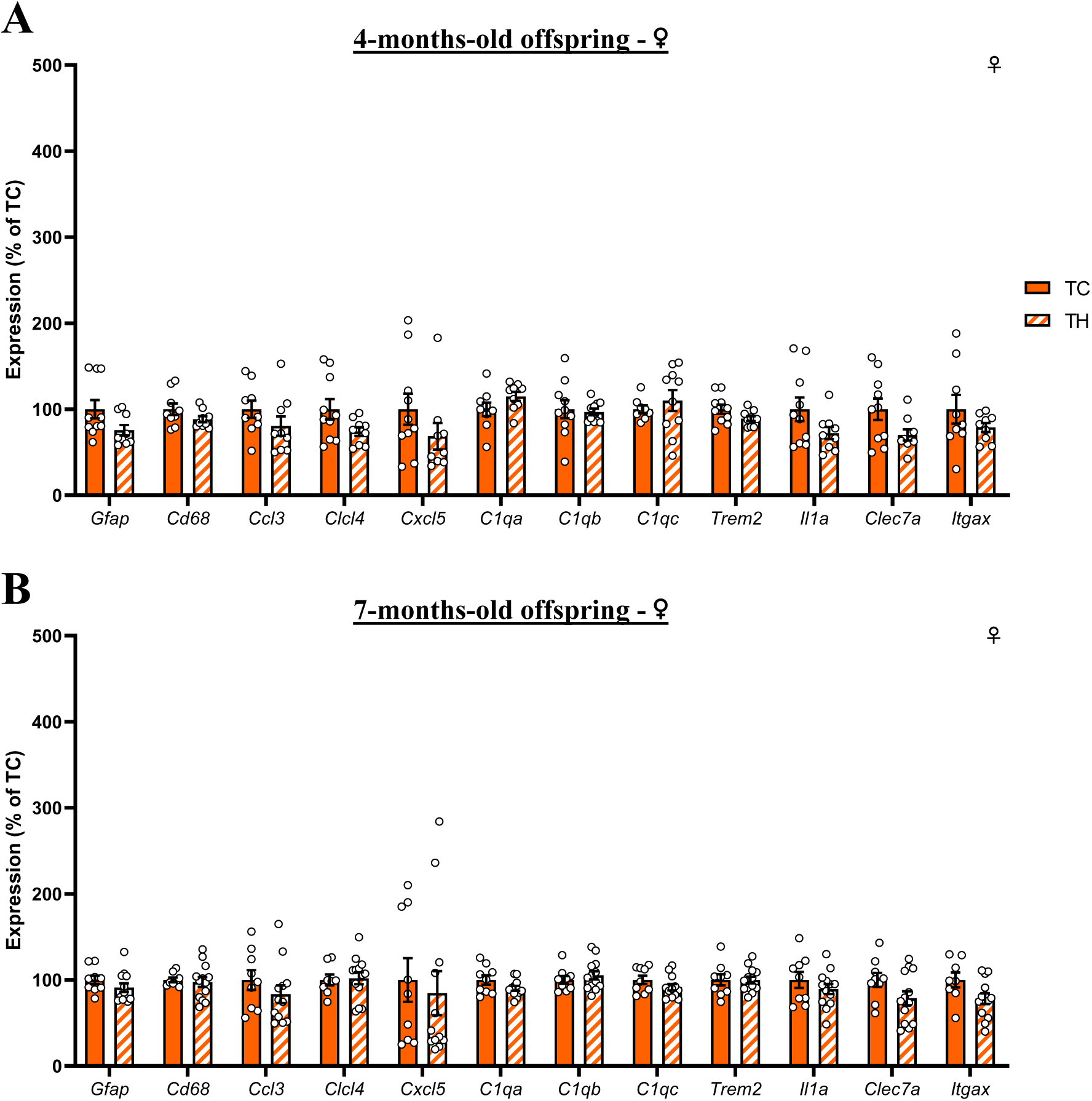
Effect of maternal HFD on hippocampal neuroinflammation in adult female THY-Tau22 offspring. (**A** and **B**) Expression of neuroinflammatory genes was measured by quantitative PCR in the hippocampus of 4-month-old and 7-month-old femaleTHY-Tau22 offspring. *n* = 7-13 per group. Values are expressed as mean ± SEM.

**Supplementary Figure 7.**
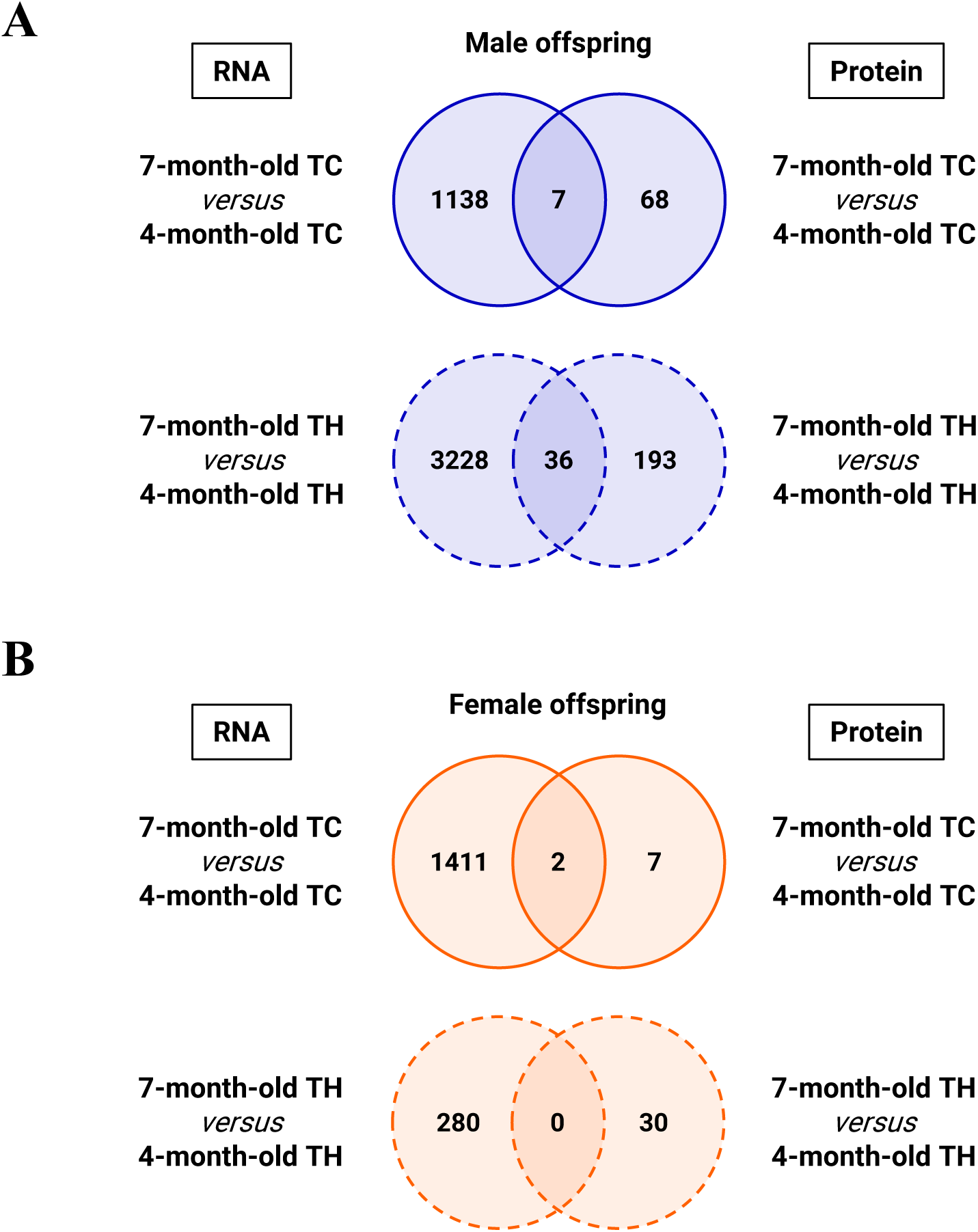
Comparison of deregulated genes and proteins between 4- and 7-months THY-Tau22 offspring.

**Supplementary Table 1.** Macronutrient and micronutrient composition of diets. Percentages of protein, fat and carbohydrate for each diet, and the associated calories in kilocalories per kilogram of body mass (kcal/kg). The sources of each macronutrient and micronutrient are indicated.

|  | Chow diet (control) | High-fat diet | Standard diet |
| --- | --- | --- | --- |
| Period | From P0 to P21 |  | From P21 to sacrifice |
| Reference | Safe A03 | Research diets D12331 | Safe A04 |
| % of proteins | 25.2 | 16.5 | 19.3 |
| Sources of proteins | Fish | Casein, di-methionin | Fish |
| % lipids | 13.5 | 58.0 | 8.4 |
| Sources of lipids | Soy extracts | Soybean oil (6%)<br>Coconut oil (94%) | Soy extracts |
| % carbohydrates | 61.3 | 25.5 | 72.4 |
| Sources of carbohydrates | Wheat, corn, barley | Maltodextrine, sucrose | Wheat, corn, barley |
| Minerals | P, Ca, Na, K, Mg, Mn,<br>Fe, Cu, Zn, Cl | P, Ca, Na, K, Mg, Mn, Fe,<br>Cu, Zn, Cl, S, Cr, I, Se | P, Ca, Na, K, Mg, Mn,<br>Fe, Cu, Zn, Cl |
| Vitamins | A, D3, B1, B2, B5, B6, B12, E, K3, niacin, folic acid, biotin, cholin |  |  |
| Kcal/Kg | 3395.0 | 5558.5 | 3339.0 |

**Supplementary Table 2.**
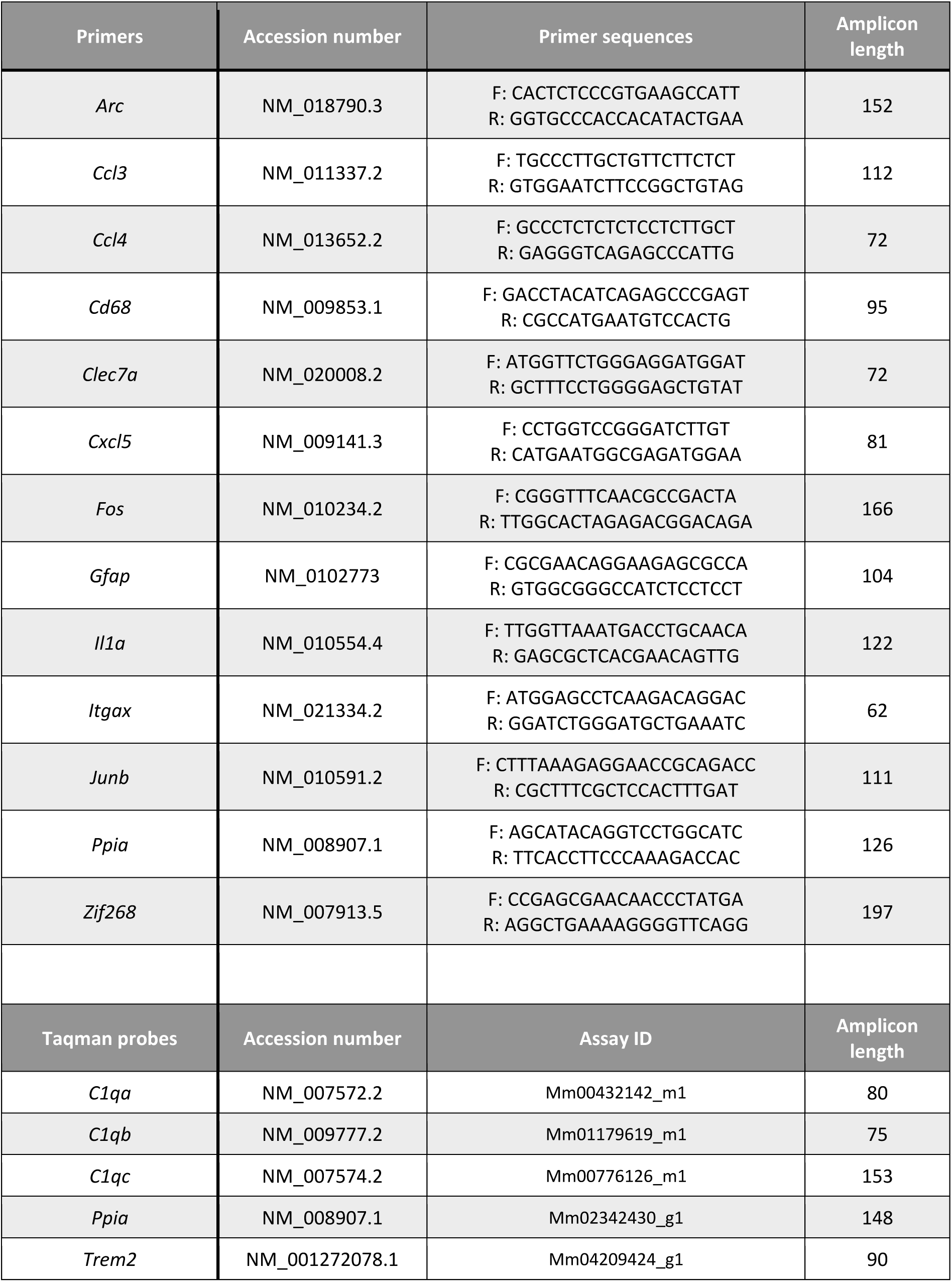
List of sequences of primers used for quantitative polymerase chain reaction analyses. F = forward; R = reverse.

**Supplementary Table 3.** List of antibodies used for western blot analysis. For homemade antibody characterizations, see Buée-Scherrer *et al.*, 1996^1^ for Tau N-terminal, Tau C-terminal and Tau pS396/S404, and Troquier *et al*., 2012^2^ for Tau pS422.

| Antibody | Species | Reference | Dilution |
| --- | --- | --- | --- |
| Tau N-terminal | Rabbit | Homemade | 1/10000 |
| Tau C-terminal | Rabbit | Homemade | 1/10000 |
| Tau pY18 | Mouse | Medimabs NM-0194-200 | 1/5000 |
| Tau pS212/T214 | Mouse | Invitrogen MN1060 | 1/1000 |
| Tau pT181 | Mouse | Invitrogen MN1050 | 1/1000 |
| Tau pT214 | Rabbit | Life Technology 44742G | 1/1000 |
| Tau pS396 | Rabbit | Invitrogen 44752G | 1/10000 |
| Tau pS396/S404 | Mouse | Homemade | 1/2 |
| Tau pS404 | Rabbit | Invitrogen A44758G | 1/10000 |
| Tau pS422 | Mouse | Homemade | 1/1000 |
| Tau1 | Mouse | Millipore MAB3420 | 1/1000 |
| NR2B | Rabbit | Millipore 06-600 | 1/1000 |
| SNAP25 | Mouse | Santa Cruz 376713 | 1/1000 |
| Beta-actin | Mouse | Sigma Aldrich A5451 | 1/10000 |
| Anti-rabbit | Goat | Vector PI-1000 | 1/5000 |
| Anti-mouse | Horse | Vector PI-2000 | 1/50000 |

**Supplementary Table 4.**
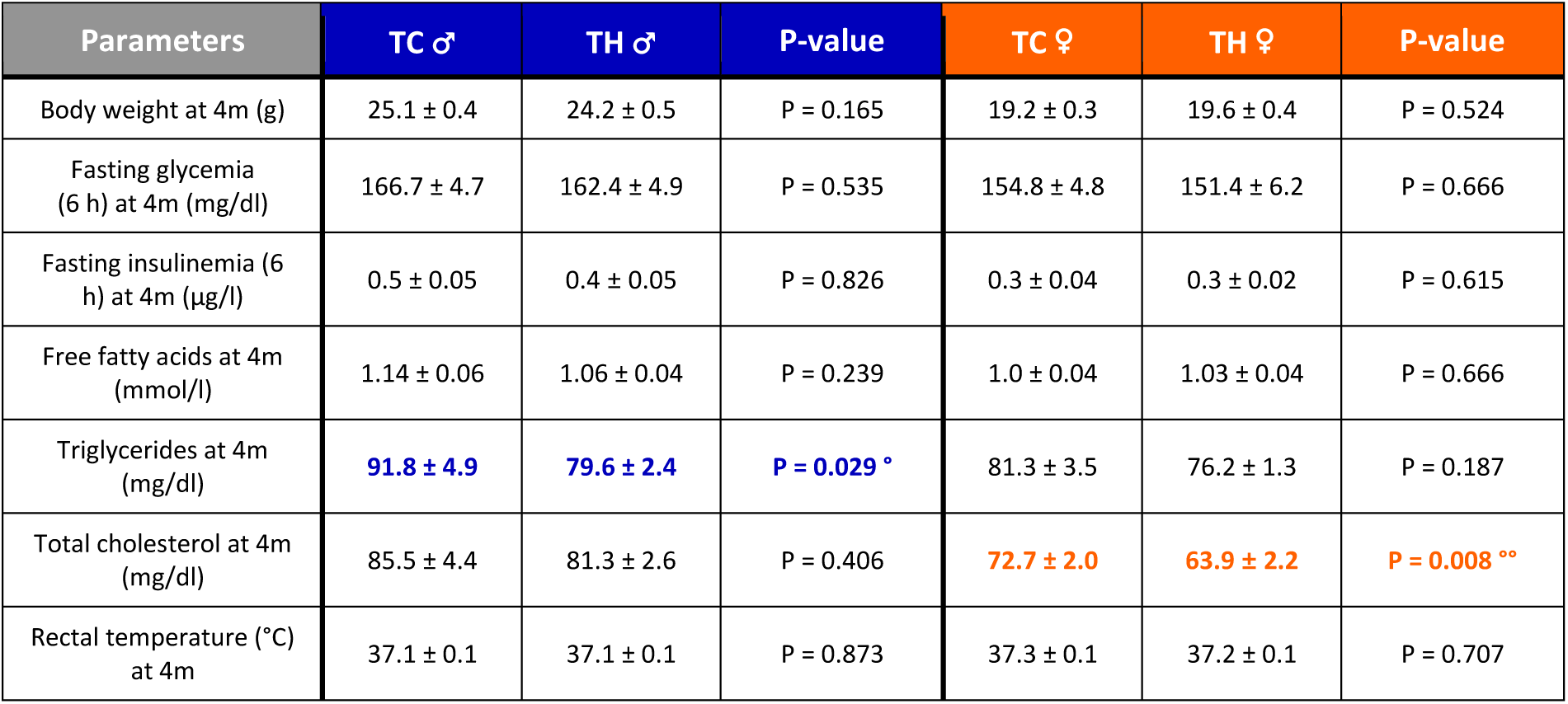
Effect of maternal HFD on physiological and metabolic parameters of 4-month-old THY-Tau22 offspring. °*P* < 0.05, °°*P* < 0.01 versus TC mice using Student’s *t*-test. *n* = 19-26 per group. Values are expressed as mean ± SEM.

| Parameters | TC ♂ | TH ♂ | P-value | TC ♀ | TH ♀ | P-value |
| --- | --- | --- | --- | --- | --- | --- |
| Body weight at 4m (g) | 25.1 ± 0.4 | 24.2 ± 0.5 | P = 0.165 | 19.2 ± 0.3 | 19.6 ± 0.4 | P = 0.524 |
| Fasting glycemia (6 h) at 4m (mg/dl) | 166.7 ± 4.7 | 162.4 ± 4.9 | P = 0.535 | 154.8 ± 4.8 | 151.4 ± 6.2 | P = 0.666 |
| Fasting insulinemia (6 h) at 4m (µg/l) | 0.5 ± 0.05 | 0.4 ± 0.05 | P = 0.826 | 0.3 ± 0.04 | 0.3 ± 0.02 | P = 0.615 |
| Free fatty acids at 4m (mmol/l) | 1.14 ± 0.06 | 1.06 ± 0.04 | P = 0.239 | 1.0 ± 0.04 | 1.03 ± 0.04 | P = 0.666 |
| Triglycerides at 4m (mg/dl) | **91.8 ± 4.9** | **79.6 ± 2.4** | **P = 0.029 °** | 81.3 ± 3.5 | 76.2 ± 1.3 | P = 0.187 |
| Total cholesterol at 4m (mg/dl) | 85.5 ± 4.4 | 81.3 ± 2.6 | P = 0.406 | **72.7 ± 2.0** | **63.9 ± 2.2** | **P = 0.008 °°** |
| Rectal temperature (°C) at 4m | 37.1 ± 0.1 | 37.1 ± 0.1 | P = 0.873 | 37.3 ± 0.1 | 37.2 ± 0.1 | P = 0.707 |

**Supplementary Table 5.**
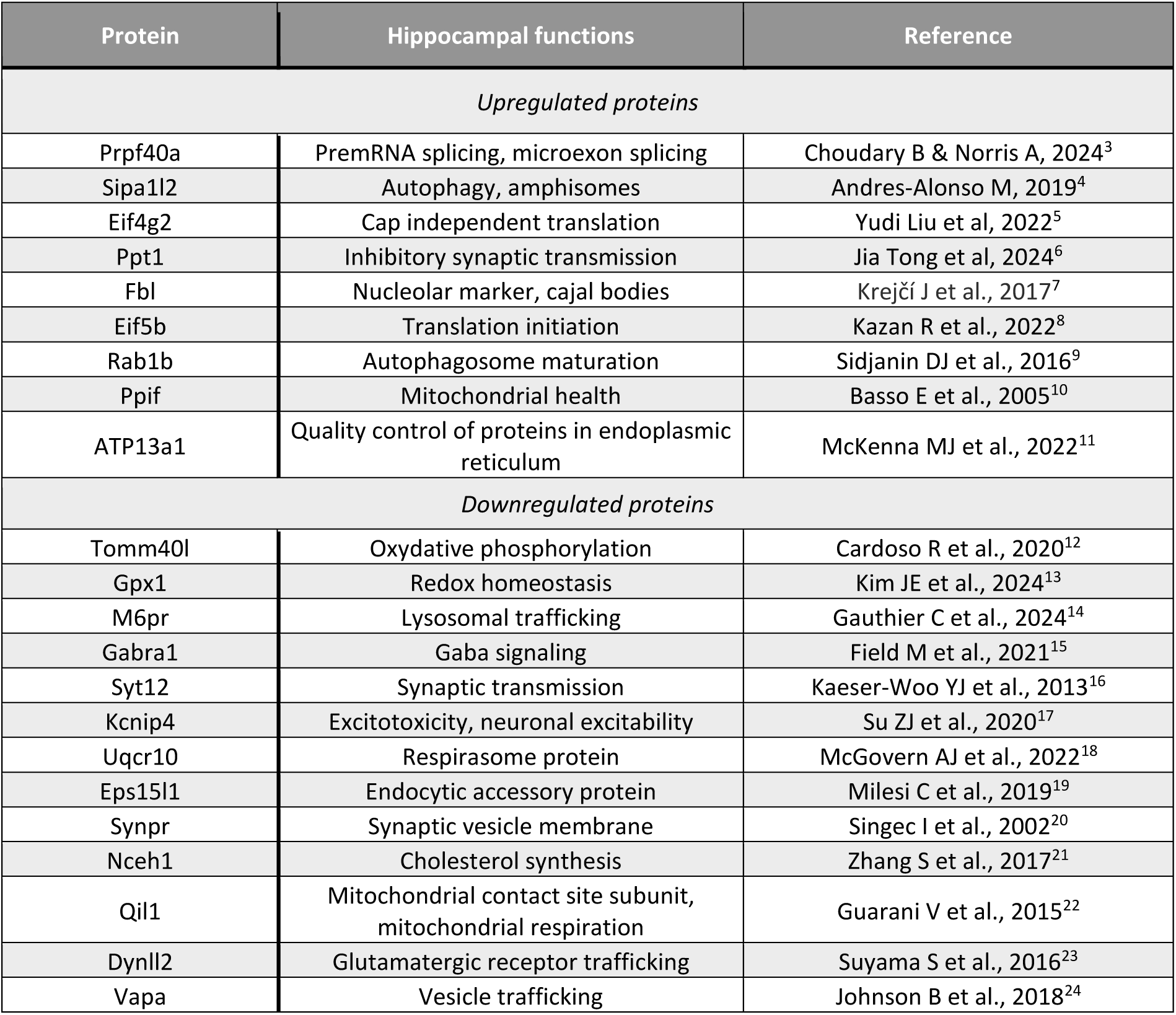
Proteins exclusively deregulated between 4-month-old and 7-month-old in male TC mice.

**Supplementary Table 6.** Proteins exclusively deregulated between 4-month-old and 7-month-old in female TC mice.

| Protein | Hippocampal functions | Reference |
| --- | --- | --- |
| *Upregulated proteins* |  |  |
| Rabgap1l | Vesicular trafficking | Li Q et al., 2022<sup>25</sup> |
| *Downregulated proteins* |  |  |
| Paip1 | Translation enhancer | Xue M et al., 2023<sup>26</sup> |
| Dkc1 | Ribosome biogenesis | Gu BW et al., 2015<sup>27</sup> |
| Itpa | Nucleotide biosynthesis | Schroader JH et al., 2023<sup>28</sup> |

**Supplementary Table 7.**
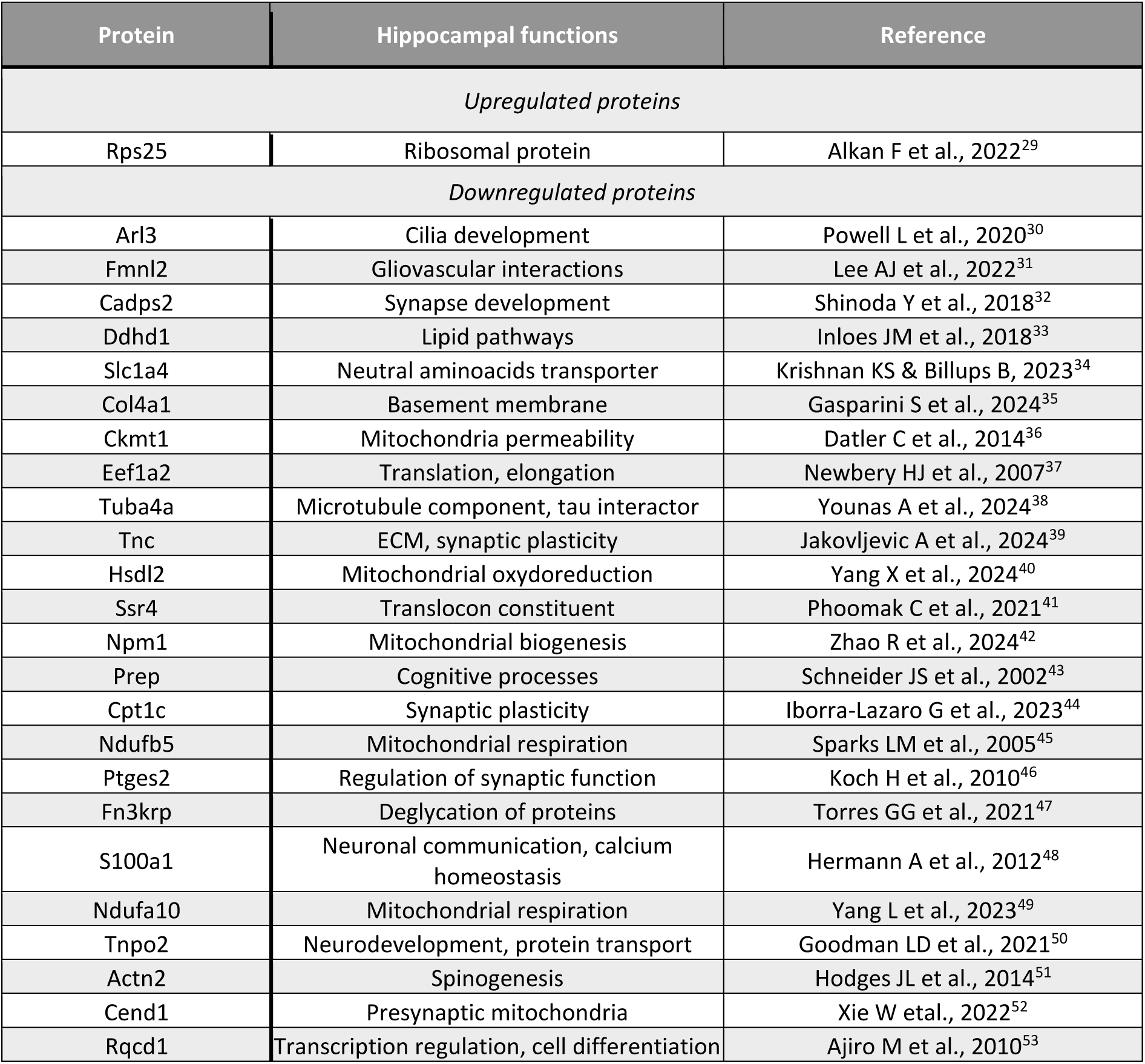
Proteins exclusively deregulated between 4-month-old and 7-month-old in female TH mice.

