## Supplementary material & methods for "Sex-dependent effects of maternal high-fat diet during lactation in adult THY-Tau22 mice offspring"

### Supplementary materials and methods

#### Behavioural studies

##### Actimetry

To investigate the spontaneous activity, mice were placed individually in the centre of an infrared actimeter (45 cm x 45 cm x 35 cm, BIOSEB LE8816), composed by a two-dimensional square frame, for 10 minutes. Spontaneous activity of mice was tracked with distance move and velocity recorded by Actitrack software (Bioseb).

##### Elevated plus maze

To investigate the anxiety-related behaviour, the elevated plus maze was used. The apparatus consists of a plus-shaped maze with two closed and two open arms (30 cm long x 6.5 cm wide) raised by 80 cm. Mice were placed at the centre of the maze with their face in the direction of a closed arm and were allowed to explore freely for 5 minutes. Time spent in open arms was recorded using EthoVision XT14 tracking system (Noldus).

##### Y-maze

Short-term spatial memory was evaluated using the Y-maze task. The apparatus consists of a Y-shaped maze with three closed arms (22 cm long, 6.4 cm wide and 15 cm high) half-filled with litter, surrounded by spatial cues. During the learning phase, mice were placed at the extremity of the “start” arm and one of the two other arms was closed (“new” arm). Mice were allowed to explore freely the “start” and “familiar” arms for 5 minutes and were put back in their home cage for 2 minutes. The position of the “familiar” and “new” arms was randomly assigned to the mice and the litter was mixed after each trial. During the retention phase, mice were placed at the extremity of the “start” arm and had freely access to the three arms for 1 minute. The time spent in each arm during the two phases was recorded by EthoVision XT14 software (Noldus). For the learning phase, the percentage of time spent in the “familiar” arm compared with the time spent in the “familiar” and “start” arms was calculated. For the test phase, the percentage of time spent in the ”new” arm. Proper short spatial memory is reflected when mice spent significantly more than 50% of time in the “new” over the “familiar arm”.

##### Barnes maze

Long-term spatial memory was evaluated using the Barnes maze task. This maze is an elevated white circular PVC open platform (80 cm height, 120 cm diameter) with 40 equally spaced holes (5 cm of diameter) located at 5 cm from its circumference and a black escape box located under one of the 40 holes. This platform is surrounded by spatial cues to help guide the animal to the escape hole. To encourage the mouse to enter the escape box, a bright light was placed over the maze (900 lux). First, for the habituation step, mice were free to explore the maze and the escape box for 3 minutes. Next, mice completed 4 days of acquisition training with four trials per day. For each trial, mice were placed in a start tube in the centre of the maze (for 5 s) and then trained to locate the escape hole (randomized for all mice) using spatial cues surrounding the maze. If a mouse did not enter the escape hole within 3 minutes it was gently guided to the escape hole. Mice remained 60 s in the escape box before returning to their home cage. The inter-trial interval was 15 minutes. After each test, the maze was cleaned with 70% ethanol to avoid odour bias, and the maze was rotated clockwise a quarter turn every day. For each trial, distance travelled was measured by EthoVision XT14 tracking system (Noldus). The retention step takes place 24 hours after the last test. Each mouse was placed on the maze without the escape box for 2 minutes. The percentage of time spent in the target quadrant i.e. the quadrant the included previously the escape box at the target hole, was calculated. In addition, the percentage of mice that explored the cumulative zone, which corresponds to the target hole and the two neighbouring holes, was also measured. Mice exploring the maze less than 30 s were excluded (one WC, one WH, four TC and two TH male mice, and two TC and one TH female mice).

#### Metabolic analysis

##### Glycemia

Blood glucose measurements in mice were performed after 6 h of fasting (postabsorptive condition) at 4 months of age. A drop of blood, obtained after incision of the tail vein, was applied to a OneTouch strep linked to a OneTouch Verio Flew glucometer.

##### Glucose tolerance test

Glucose tolerance was assessed at 3 months of age using the intraperitoneal glucose tolerance test (IPGTT) following 6 h of morning fasting. D (+) glucose (1 g/kg; Sigma-Aldrich G8270) was injected intraperitoneally. Blood glucose was then measured (see section 4.a for the protocol) at 0, 15, 30, 60, 90 and 120 minutes following injection.

##### Biochemical plasma parameters

Blood was collected at the tail vein after 6 hours of fasting in 4-month-old animals. Blood was centrifuged at 1,500 g for 15 minutes at 4°C and plasma was separated and transferred in a new tube. Plasma concentration of insulin was measured using the mouse insulin ELISA kit (Mercodia AB 10-1247-01; no cross-reactivity with proinsulin) following the manufacturer's instructions. Commercially available kits were used to measure total cholesterol (Thermo Fisher Scientific MG981813), triglycerides (Diasys 157109910026) and free fatty acids (Diasys 157819910935) plasma concentrations applied to a biochemistry analyzer (Thermo Fisher Scientific Konelab20) with colorimetric methods.

#### Quantitative polymerase chain reaction

500 ng of total RNA was reverse-transcribed using the high-capacity cDNA reverse transcription kit (Applied Biosystems 4368814). Quantitative real-time reverse transcription-PCR analysis was performed on a StepOnePlus system (Applied Biosystems) using Power SYBR Green PCR master mix (Applied Biosystems 4367559). The thermal cycler conditions were as follows: 2 minutes at 50°C, 10 minutes at 95°C, 40 cycles of a two-step PCR consisting of a 95°C step for 15 s followed by a 60°C step for 25 s, 15 s at 95°C, followed by a melting curve protocol. Sequences of primer used are given in the Supplementary Table 2. *Peptidylprolyl isomerase A* (*Ppia*) was used as internal control. Amplifications were carried out in duplicate, and the relative expression of target genes was determined by the delta-delta-cycle threshold method.

#### Biochemical analysis

##### Protein extraction

Hippocampal frozen tissue was sonicated on ice in 200 µl of Tris buffer containing 10% sucrose (pH 7.4), homogenized at 4°C for 1 hour using a rotating mixer and stored at -80°C until western blot or mass spectrometry analysis.

##### Synaptosomes extraction

Hippocampal frozen tissue was homogenized using a potter on ice in 200 µl of Tris buffer containing 10% sucrose (pH 7.4). 400 µg of proteins were diluted in 400 µl of synaptic protein extraction buffer (Thermo Fisher Scientific 87793) and then homogenized using a potter. After a first centrifugation (1,200 g for 10 minutes), the supernatant was collected to do a second centrifugation (15,000 g for 20 minutes). Finally, the pellets corresponding to the synaptosomes fraction were diluted in 100 µl of synaptic protein extraction buffer.

##### Western blot

Protein concentrations were quantified using a bicinchoninic acid (BCA) protein assay (Thermo Fisher Scientific 1023539), subsequently diluted with lithium dodecyl sulphate (Invitrogen NP0009) buffer supplemented with reducing agents (Invitrogen NP0007), and then separated on 4-12% Bis-Tris polyacrylamide gels. Proteins were transferred to nitrocellulose membranes, which were then saturated by 5% non-fat dry milk, or 5% bovine serum albumin diluted in Tris 15 mM, NaCl 140 mM and Tween 20 0.1% buffer (pH 8) and incubated with the primary antibody for 24 hours at 4°C and with the appropriate secondary antibody for 1 hour at room temperature (RT). Antibodies used are given in Supplementary Table 3. Signals were visualized using a chemiluminescence kit (Amersham RPN2106) and an Imager 800. Results were normalized to β-actin and quantification was performed using ImageJ software. After this first normalization, the phosphoepitopes of tau protein were normalized to total tau protein.

#### Immunostaining analysis

##### Synaptic proteins

Brains from 7-month-old mice were cut to obtain 35 µm-thick coronal floating sections using a cryostat. Tissue was permeabilized with 0.2% Triton XT-100 three times for 10 minutes. Nonspecific binding was blocked by the incubation with an appropriate blocking solution (mouse-on-mouse solution, Vector Laboratories MKB-2213; normal goat serum, Vector Laboratories S1000) for 1 hour at RT. Sections were then incubated at 4°C for one day with anti-PSD95 (dilution 1:200, Thermo Fisher Scientific 51-6900) or anti-SYP (dilution 1:1,000, Abcam ab14692). After washing with phosphate-buffered saline (PBS), sections were incubated with Alexa Fluor 568 nm goat anti-rabbit (dilution 1:500, Invitrogen A11011) for 1 hour at RT. Cell nuclei were stained with 4',6-diamidino-2-phenylidol (DAPI, Thermo Fisher

Scientific 62248) for 5 minutes and sections were incubated in Sudan Black buffer (Merck Millipore 2160) for 5 minutes to reduce tissue autofluorescence.

#### EDU/DCX and EDU/NeuN co-labelling

EDU/Doublecortin (DCX) and EDU/Neuronal nuclear antigen (NeuN) co-labelling was performed on 35 µm-thick coronal floating brain sections obtained using a cryostat. Each labelling was performed on two sections per mouse, one containing the anterior (bregma -1.82) and one the posterior (bregma -2.54) hippocampus. For EDU/DCX co-labelling, immunofluorescence with rabbit anti-DCX antibody (1:500 dilution, Abcam ab2253) followed the same protocol as presented in section 6.e.i. However, after incubation with DAPI, EDU was revealed following the protocol outlined in section 5.b. EDU/NeuN co-labelling is similar with one exception: the primary anti-NeuN antibody made in mice was biotinylated (1:500 dilution, Merck Millipore MAB337B), while the anti-mouse secondary antibody was linked to a streptavidin coupled to a fluorophore emitting at 568 nm (Invitrogen S11226).

#### Tri-dimensional analysis of EDU/DCX cells

The tri-dimensional analysis of EDU/DCX cells was performed on EDU/DCX co-labelling on 50 µm-thick coronal floating brain sections obtained using a vibratome. Each labelling was performed on serial coronal sections representing the entire hippocampus. The protocol for EDU/DCX co-labelling is the same as previously mentioned. However, tissue was permeabilized with 0.2% Triton XT-100 three times for 1 day, the rabbit anti-DCX primary antibody was incubated for 3 days, and the anti-mouse secondary antibody was incubated for 1 day.
